# Behavioral and genomic sensory adaptations underlying the pest activity of *Drosophila suzukii*

**DOI:** 10.1101/2020.10.15.341594

**Authors:** Sylvia M. Durkin, Mahul Chakraborty, Antoine Abrieux, Kyle M. Lewald, Alice Gadau, Nicolas Svetec, Junhui Peng, Miriam Kopyto, Christopher B. Langer, Joanna C. Chiu, J.J. Emerson, Li Zhao

## Abstract

Studying how novel phenotypes originate and evolve is fundamental to the field of evolutionary biology as it allows us to understand how organismal diversity is generated and maintained. However, determining the basis of novel phenotypes is challenging as it involves orchestrated changes at multiple biological levels. Here, we aim to overcome this challenge by using a comparative species framework combining behavioral, gene expression, and genomic analyses to understand the evolutionary novel egg-laying substrate-choice behavior of the invasive pest species *Drosophila suzukii*. First, we used egg-laying behavioral assays to understand the evolution of ripe fruit oviposition preference in *D. suzukii* as compared to closely related species *D. subpulchrella* and *D. biarmipes*, as well as *D. melanogaster*. We show that *D. subpulchrella* and *D. biarmipes* lay eggs on both ripe and rotten fruits, suggesting that the transition to ripe fruit preference was gradual. Secondly, using two-choice oviposition assays, we studied how *D. suzukii, D. subpulchrella, D. biarmipes* and *D. melanogaster* differentially process key sensory cues distinguishing ripe from rotten fruit during egg-laying. We found that *D. suzukii*’s preference for ripe fruit is in part mediated through a species-specific preference for stiff substrates. Lastly, we sequenced and annotated a high-quality genome for *D. subpulchrella*. Using comparative genomic approaches, we identified candidate genes involved in *D. suzukii*’s ability to seek out and target ripe fruits. Our results provide detail to the stepwise evolution of pest activity in *D. suzukii*, indicating important cues used by this species when finding a host, and the molecular mechanisms potentially underlying their adaptation to a new ecological niche.

## INTRODUCTION

Novel phenotypes can give species the opportunity to occupy a new ecological niche (Mayr, 1960; Moczek, 2008; Muller and Wagner, 1991). Understanding how and when these phenotypes arise is an exciting question in evolutionary biology. Adaptive traits can present as changes to an organism’s behavior, physiology, or morphology, and arise through a variety of different molecular mechanisms. In the context of pest species, adaptation to a new ecological niche can come with damaging environmental, agricultural, and economic consequences. Understanding the basis of novel pest behavior can also shed light on the ecological impact of invasive species. Unlike the majority of *Drosophila* flies, the species *Drosophila suzukii* prefers to lay its eggs in ripe as opposed to rotten fruit, causing substantial crop damage and leading to economic losses in the fruit industry (Rota-Stabelli et al., 2013). Although originally categorized in Japan and likely to be native to East Asia, within the past decade *D. suzukii* has spread throughout Europe and North America (Adrion et al., 2014; Fraimout et al., 2017; Paris et al., 2020). Preference for ripe fruit is thought to have evolved along the lineage leading to *D. suzukii*, as *D. melanogaster* and other outgroup species, including *D. takahashii, D. eugracilis*, and *D. ananassae*, all have an oviposition preference for rotten fruit (Karageorgi et al., 2017). *D. suzukii* has both physical and behavioral traits that allow it to selectively target ripe fruit as an oviposition substrate (Atallah et al., 2014). Morphologically, *D. suzukii* has evolved an enlarged, serrated ovipositor allowing females to puncture hard surfaces and insert their eggs into ripe fruit. Behaviorally, they have evolved the ability to seek out and selectively target ripe fruits for oviposition through changes in multiple sensory systems (Karageorgi et al., 2017). Thus, the exploitation of the ripe fruit niche by *D. suzukii* requires orchestrated changes at multiple biological levels. Investigating the behavioral and molecular underpinnings of these changes will advance our understanding of the forces and mechanisms that drove *D. suzukii*’s evolution into an invasive pest.

Adaptive shifts in host preference, such as that described for *D. suzukii*, are often mediated through sensory system evolution. Changes in the function and sequence of chemoreceptor genes – including odorant receptors (ORs), ionotropic receptors (IRs), and gustatory receptors (GRs) – and mechanosensory receptor genes (MRs) underlie various species-specific host preference differences in insects (Auer et al., 2020; Dekker et al., 2006; Goldman-Huertas et al., 2015; Karageorgi et al., 2017; Mansourian et al., 2018; Vosshall and Stocker, 2007). For example, modifications to sensory receptor genes are linked to the transition to herbivory in *Scaptomyza flava* (Goldman-Huertas et al., 2015), the oviposition preference for morinda fruits (*Morinda citrafolia*) in *D. sechellia* (Auer et al., 2020; Dekker et al., 2006), and the specialization on marula fruit in ancient *D. melanogaster* (Mansourian et al., 2018). Additionally, through knockout experiments in *D. melanogaster*, sensory gene function has been linked to various behaviors relevant to *D. suzukii* pest activity, including preference of acetic acid containing oviposition sites (Joseph et al., 2009), the avoidance of stiff substrates (Jeong et al., 2016; Zhang et al., 2020, 2016), and the detection of substances that are at different concentrations in ripe and rotten fruit – such as CO_2_, acetic acid, and sugar (Fujii et al., 2015; Kwon et al., 2007; Rimal et al., 2019).

While progress has been made in understanding the evolution of olfactory genes in *D. suzukii* (Keesey et al., 2015; Ramasamy et al., 2016), very little is known about the evolutionary history of other sensory genes in this system, despite *D. suzukii*’s oviposition site preference being mediated via multiple sensory systems (Karageorgi et al., 2017). Finally, studies of adaptive sensory gene evolution in *D. suzukii* often focus only on changes to the coding sequence of these genes, despite differential gene expression playing a prominent role in the evolution of adaptive phenotypes (Carroll, 2005; Jones et al., 2012; Phifer-Rixey et al., 2018; Wray, 2007). For example, changes in expression of ORs and odorant-binding proteins in the olfactory organs of *D. sechellia* are thought to in part mediate this species’ specialization on morinda fruits (Kopp et al., 2008).

Although the phenotypic and behavioral innovations leading to the pest status of *D. suzukii* are overall well defined, the specific sensory cues used by *D. suzukii* to target ripe fruits as well as the molecular changes accompanying these sensory changes remain unknown. Here we examine these knowledge gaps by investigating the behavioral, genomic, and gene regulatory mechanisms underlying the pest activity of *D. suzukii*.

First, we provide new insights into the stepwise evolution of ripe fruit preference in *D. suzukii* through the in-depth behavioral and genomic examination of two closely related species, *D. subpulchrella* and *D. biarmipes*. Second, we identified key sensory cues of ripe and rotten fruit differentially processed by *D. suzukii, D. subpulchrella, D. biarmipes* and *D. melanogaster* in the context of egg-laying preference. Third, to infer the most recent changes in genome content in *D. suzukii*, we sequenced the genome of the closely related species, *D. subpulchrella*, and performed comparative genomic analyses. Lastly, we identified candidate genes involved in *D. suzukii*’s ability to seek out and target ripe fruits through population level analyses of olfactory, gustatory, and mechanosensory receptor genes with differential expression and signatures of positive selection in *D. suzukii* as compared to *D. subpulchrella, D. biarmipes* and *D. melanogaster*. We provide a view, from multiple levels of analysis, of the origin of pest activity in *D. suzukii*, indicating important sensory cues used by this species during egg-laying, and the gene expression and genomic changes potentially underlying this novel pest behavior.

## RESULTS AND DISCUSSION

### Ripe fruit preference in *D. suzukii* evolved after the split with *D. subpulchrella*

To clarify the evolutionary history of *D. suzukii*’s oviposition behavior, we built upon previous work that established a step wise evolution of ripe fruit oviposition preference in *D. suzukii*, with *D. melanogaster* preferring rotten fruit and *D. biarmipes* showing an equal preference for ripe and rotten fruit (Karageorgi et al., 2017) by measuring the oviposition preference of *D. suzukii*’s sister species, *D. subpulchrella*. It was previously hypothesized that *D. subpulchrella* prefers ripe fruit for oviposition, as *D. suzukii* does (Karageorgi et al., 2017), because they share the trait of an enlarged ovipositor and the ability to lay eggs in ripe fruits (Atallah et al., 2014). However, this assumption has not been empirically tested before this study. To test this hypothesis, we placed female flies in a cage with both a ripe and rotten whole strawberry and allowed them to lay eggs overnight. Afterwards, the number of eggs laid in each fruit was counted and an oviposition preference index (OPI) was calculated (Figure 1A, supplementary file 1).

**Figure 1.**
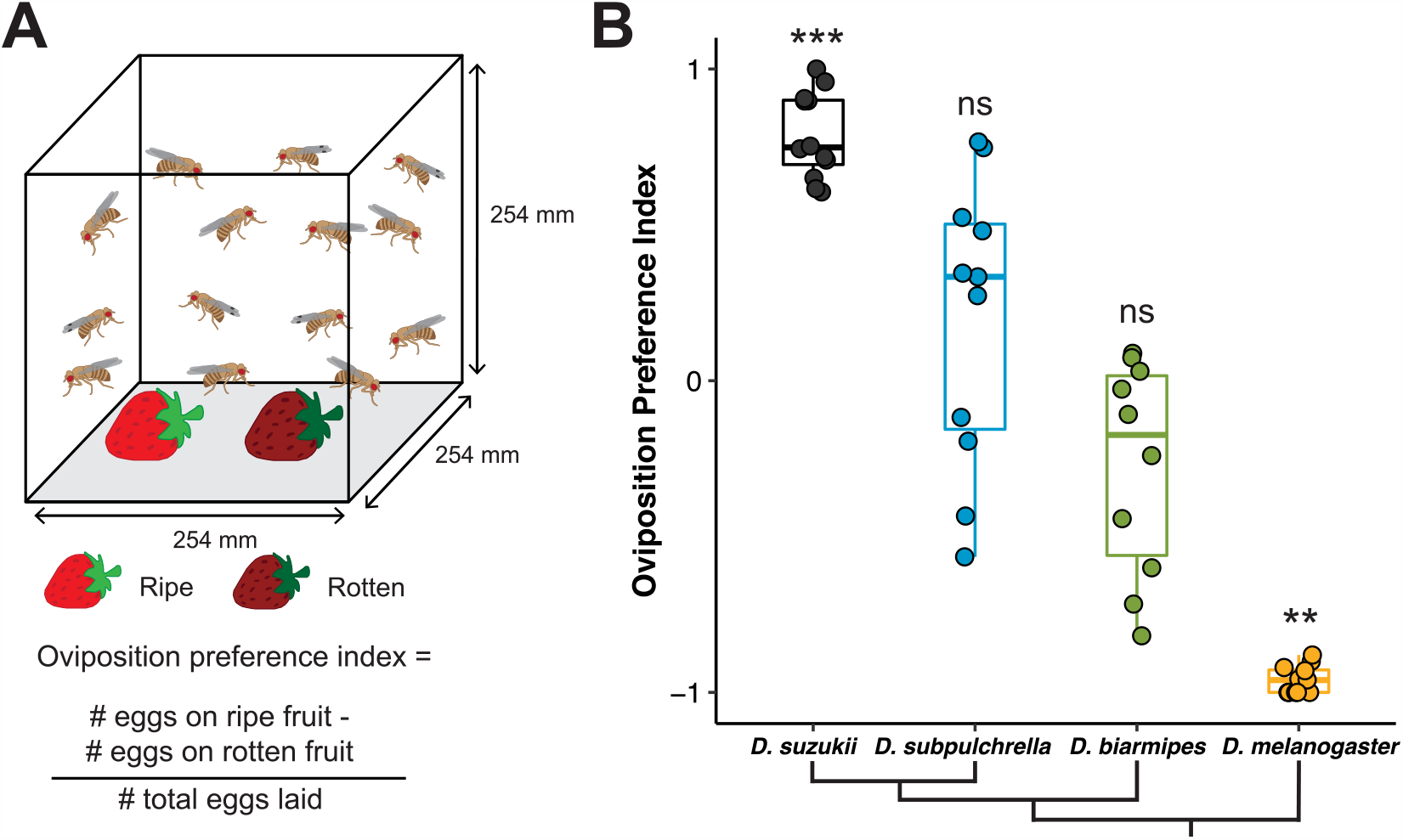
The evolution of ripe fruit preference in *D. suzukii* occurred gradually, with *D. subpulchrella* and *D. biarmipes* as intermediates. A. Schematic of two-choice oviposition preference assay. 10 females and 5 males were placed in a cage with a ripe and rotten strawberry for 19 hours. Number of eggs on each fruit was then counted. B. Oviposition preference index of four focal species. *D. suzukii* prefers to oviposit in ripe fruit, *D. subpulchrella* and *D. biarmipes* show no preference for either fruit, and *D. melanogaster* prefers rotten fruit. Each data point represents one experimental trial and data dispersion is represented by a boxplot. Preference P-values were calculated using a Wilcoxon signed rank test against a theoretical value of 0 (no preference). ***, p≤0.001; **, p≤=0.01; *, p≤0.05; ns, p>0.05 for all future figures.

We recapitulated the oviposition preference results for *D. suzukii, D. biarmipes*, and *D, melanogaster*, and found that, contrary to the above hypothesis, *D. subpulchrella* has no distinct preference for ripe or rotten fruit, displaying an intermediate oviposition behavior similar to *D. biarmipes* (OPI = 0.196 ± 0.138SEM) (Figure 1B).

To investigate the relative roles of stiffness and chemosensory differentiation in driving the intermediate preference of *D. subpulchrella*, we measured the oviposition preference of a cut-ripe vs. rotten strawberry for both *D. suzukii* and *D. subpulchrella* (Figure S1A). While both species can puncture and lay eggs in ripe fruit, *D. suzukii* can utilize a wider range of stiff egg-laying substrates as demonstrated by their ability to pierce the skin of grapes, which *D. subpulchrella* cannot do (Atallah et al., 2014). It was previously established that *D. biarmipes* and *D. melanogaster* cannot puncture ripe fruits, so we did not measure them here (Karageorgi et al., 2017). If *D. subpulchrella* oviposition site choice is not based on stiffness, we would expect a similar result to our whole fruit assay, in which *D. subpulchrella* equally prefers cut-ripe and rotten fruit. We found that *D. suzukii* still preferred the ripe fruit (Wilcoxon signed rank test, P =0.0004), while *D. subpulchrella* displayed no significant preference for the ripe or rotten fruit (Wilcoxon signed rank test, P=0.06) (Figure S1B). However, *D. subpulchrella* is trending toward having a ripe fruit preference; the mean OPI for *D. subpulchrella* in this assay (OPI = 0.341 ± 0.166SEM) was higher as compared to their whole fruit OPI (OPI = 0.196 ± 0.138SEM), and *D. suzukii* and *D. subpulchrella* preference distributions were not significantly different from one another (pairwise Wilcoxon Test, P=0.27). This suggests that both stiffness and chemosensory differentiation are involved in *D. subpulchrella* oviposition site choice. Additionally, both *D. suzukii* and *D. subpulchrella* strongly disfavor laying eggs on the exposed, white flesh of the cut fruit, further suggesting that texture differentiation is an important facet of oviposition site choice in these species (Figure S1C).

*D. subpulchrella*’s intermediate preference suggests that ripe fruit preference in *D. suzukii* is due to factors beyond the enlargement of the ovipositor and ability to puncture the skin of ripe fruit and evolved after the split between *D. subpulchrella* and *D. suzukii*. Further, because *D. suzukii* is an invasive pest while *D. subpulchrella* is not, differences between these sister species may point to traits that have contributed to the evolution of pest behavior in *D. suzukii*, and may be important for future efforts in mitigating damage from this invasive pest.

### *D. suzukii* displays a preference for stiff oviposition substrates, an aversion to acetic acid, and lack of preference for sucrose

Previous studies indicate that *D. suzukii*’s preference for ripe fruit evolved through changes in multiple sensory systems, including the olfactory, gustatory, and mechanosensory systems (Karageorgi et al., 2017; Keesey et al., 2015; Ramasamy et al., 2016). However, the evolutionary history of these sensory changes and the specific sensory cues involved in *D. suzukii*’s oviposition site choice remain unknown. To investigate these questions, we tested the oviposition preference of *D. suzukii, D. subpulchrella, D. biarmipes* and *D. melanogaster* females for four sensory cues that change over the course of fruit maturation from ripening to rotting: ethanol, sucrose, acetic acid, and stiffness (using agarose concentration as a proxy). At early pre-ripe and ripe maturation stages, fruit is firm and contains low levels of ethanol and total acid, both ranging from concentrations of 0 to 1% in strawberries and other fruits (Dudley, 2004; Hidalgo et al., 2012; Montero et al., 1996). In rotten fruit, as fermentation and acidification occur, ethanol and acid concentrations both rise to about 7% for ethanol (Dudley, 2004; Hidalgo et al., 2012), and about 3.5% for acetic acid (Hidalgo et al., 2012). Sugar content is highest in ripe fruit, and differs greatly in various fruit species ranging from concentrations of about 7% to 20% (translating to 200mM to 600mM sucrose) (Basson et al., 2010; Dudley, 2004; Hidalgo et al., 2012; Montero et al., 1996). We tested oviposition preference for each sensory cue at three biologically relevant levels in the context of fruit maturation (see methods) by placing 3-4 females of each species in custom built egg-laying chambers and giving them the choice between two agarose egg-laying substrates: one control and one containing the experimental substrate (Figure 2A, supplementary file 2). Afterwards, the number of eggs laid on each substrate was counted and the OPI was calculated.

**Figure 2.**
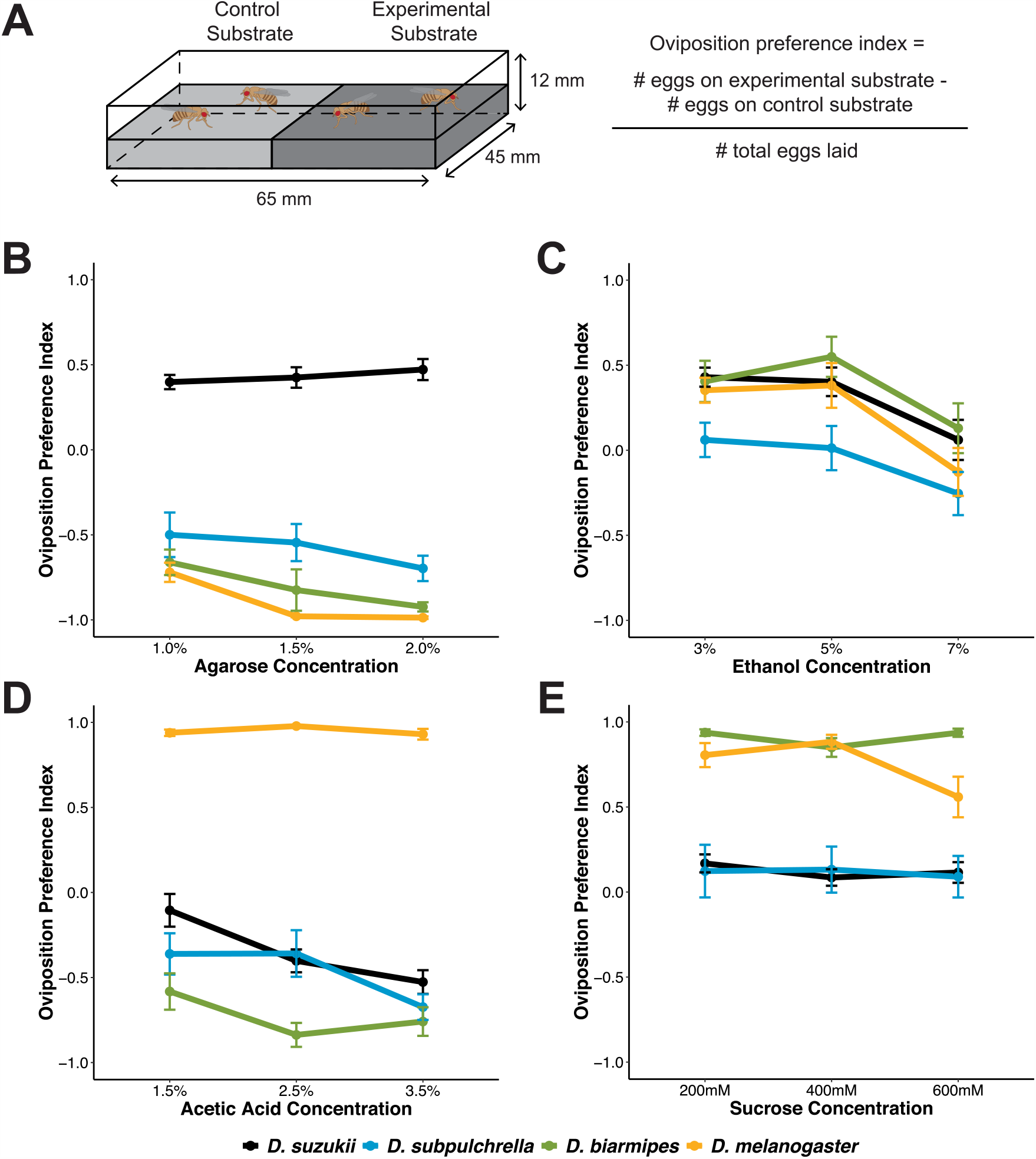
Oviposition preference for substrates associated with fruit maturation differ among focal species. A. Schematic of egg laying chamber for the substrate gradient two-choice oviposition preference assay. 3-4 females were placed in an arena with a choice between two substrates for 19 hours. Number of eggs on each substrate was then counted. Experimental substrate was varying concentrations of either ethanol, sucrose, acetic acid, or agarose. n ≥ 15 for each species at each concentration. P-values calculated through linear regression analysis through pairwise comparisons of the y-intercepts of each species’ preference curve across the three concentrations (see methods). Data points are mean ± SEM. B. Stiffness preference separated *D. suzukii* from the other three species, as it was the only species that consistently prefers stiffer oviposition substrates. There were significant differences in the preference curves of *D. suzukii* and *D. subpulchrella* (*), *D. suzukii* and *D. biarmipes* (**), *and D. suzukii* and *D. melanogaster* (***). C. Ethanol oviposition preference did not differ between *D. suzukii, D. biarmipes*, or *D. melanogaster*, with all species showing a preference for ethanol at 3% and 5%, and a neutral response to ethanol at 7%. *D. subpulchrella* displayed a neutral response to ethanol at each concentration measured. There were no significant differences in preference curves between any two species. D. *D. melanogaster* had a strong oviposition preference for acetic acid at each concentration measured, while *D. biarmipes, D. subpulchrella*, and *D. suzukii* had an aversion to acetic acid containing substrates. There were significant differences in the preference curves of *D. suzukii* and *D. biarmipes* (**), *D. suzukii* and *D. melanogaster* (***), *D. subpulchrella* and *D. melanogaster* (***), and *D. biarmipes* and *D. melanogaster* (***). E. *D. suzukii* and *D. subpulchrella* did not show an aversion to or preference for sucrose at any concentration measured, while *D. biarmipes* and *D. melanogaster* prefer sucrose containing substrates. There were significant differences in the preference curves of *D. suzukii* and *D. biarmipes* (***), *D. suzukii* and *D. melanogaster* (***), *D. subpulchrella* and *D. biarmipes* (**), and *D. subpulchrella* and *D. melanogaster* (**).

When given the choice between 0.25% and 1%, 1.5%, or 2% agarose, *D. suzukii* consistently preferred the stiffer substrate (representative of early fruit maturation stages), while *D. subpulchrella, D. biarmipes* and *D. melanogaster* preferred the softer substrate (Figure 2B, S2A, supplementary file 2, Wilcoxon signed rank test, all P-values <0.01). *D. suzukii*’s striking divergence in stiff substrate preference suggests that the species-specific targeting of stiff oviposition sites is associated with the early fruit stage preference only exhibited by *D. suzukii*. Further, the contrasting stiffness preference of *D. subpulchrella* and *D. suzukii* may explain why *D. suzukii* is able to target a broader range of ripe, soft-skinned fruit despite both species being able to puncture ripe fruits with their enlarged ovipositors (Atallah et al., 2014).

We next measured the oviposition preference for acetic acid and ethanol, which are both products of fermentation that increase as fruit rots, and are attractive oviposition cues for *D. melanogaster* (Azanchi et al., 2013; Dudley, 2004; Eisses, 1997; Joseph et al., 2009; Jouandet and Gallio, 2015; Kacsoh et al., 2013). Using a linear regression approach comparing the difference in preference curve y-intercepts, we found that overall oviposition preference for ethanol did not differ between the four focal species (Figure 2C, see methods for details). This lack of behavioral difference despite clear differences in ripe vs. rotten fruit oviposition preference may be due to the key role of yeast, which converts fruit sugars to ethanol during fermentation, in guiding natural oviposition preference. Both *D. suzukii* and *D. melanogaster* display yeast-mediated oviposition preference, with yeast being a major attractant, more so than fruit volatiles, for *D. melanogaster* (Becher et al., 2012), and *D. suzukii* exhibiting competition dependent yeast attraction, choosing to oviposit in substrates without yeast only when other species are present (Kidera and Takahashi, 2020).

*D. melanogaster* showed a strong preference for acetic acid at each concentration tested (Wilcoxon signed rank test, all P-values <0.001), whereas *D. suzukii, D. subpulchrella*, and *D. biarmipes* avoided acetic acid at high concentrations, which are representative of late fruit maturation stages (Figure 2D, S2C, supplementary file 2, Wilcoxon signed rank test at 3.5% acetic acid, all P-values <0.01). The preference for acetic acid potentially represents a unique shift in preference for *D. melanogaster*, as opposed to a loss of preference in the *D. suzukii* lineage, as other species in the *D. melanogaster* subgroup, including *D. yakuba, D. simulans* and *D. mauritiana*, have a decreased tolerance of acetic acid (McKenzie and McKechnie, 1979; Montooth et al., 2006). Investigating the acetic acid oviposition preference of more species within and outside of the *D. melanogaster* subgroup may help elucidate the role of acetic acid avoidance in conferring ripe vs. rotten fruit preference.

Lastly, we tested the oviposition preference for sucrose in each of the four species. We found that while *D. melanogaster* and *D. biarmipes* have a preference for sucrose at each concentration measured (Wilcoxon signed rank test, all P-values <0.01), *D. suzukii* and *D. subpulchrella* show no preference or aversion to sucrose, with *D. suzukii* having no preference at each concentration measured (Wilcoxon signed rank test, all P-values >0.05), and *D. subpulchrella* having no preference at 400mM and 600mM sucrose (Wilcoxon signed rank test, all P-values >0.05) (Figure 2E, S2D, supplementary file 2). Because sugar increases during ripening, high sugar content indicates that rotting will begin imminently, perhaps explaining why *D. melanogaster* chooses high sugar substrates despite the fact that their preferred substrate of rotten fruit contains less sugar than ripe fruit. Similarly, *D. suzukii*’s indifference towards sucrose may be associated with their transition towards ovipositing on early fruit maturation stages. While in a two choice scenario *D. suzukii* prefers ripe fruit (higher sugar) over rotten fruit (lower sugar), when given the option of earlier maturation stages, they equally target pre-ripe and ripe fruits for egg-laying (Karageorgi et al., 2017). Sugar is still relatively low during pre-ripe stages (Basson et al., 2010; Montero et al., 1996), suggesting that the loss of sucrose preference in *D. suzukii* could be associated with the selection of early fruit stage oviposition sites. It was previously reported that the chemical composition and contrasting stiffness of ripe and rotten fruit together explain the difference in oviposition preference in *D. suzukii, D. biarmipes*, and *D. melanogaster* (Karageorgi et al., 2017). Here, we have identified discrete differences in the roles of sensory cues associated with the fermentation and acetification process of fruit rotting in guiding egg-laying decisions in *D. suzukii* as compared to closely related, non-pest species, *D. subpulchrella* and *D. biarmipes*, as well as *D. melanogaster*.

### Sequencing, assembly, and annotation of the *D. subpulchrella* genome

The results from our phenotypic assays suggesting strongly anchored behavioral differences between species motivated a deeper study of the genetic factors underlying these behavioral differences. In order to apply comparative genomic and genetic approaches, we generated a near-chromosomal level assembly of the *D. subpulchrella* genome using PacBio sequencing (see methods). The genome size is about 265 megabases (Mb), and has a contig N50 of 11.59Mb. Specifically, the longest 6 contigs are 29.67 Mb, 29.50 Mb, 26.21 Mb, 22.17 Mb, 20.23 Mb, and 11.59 Mb, accounting to a total of 139.38 Mb (for details see Table S1). We assessed the gene content of the *D. subpulchrella* genome and found that 98.11% (2746 out of 2799) of the Diptera BUSCO genes are present in the genome, among which, 97.43% (2727) are complete and single-copy genes. When using the 303 BUSCO eukaryota genes, 302 complete genes were found in the genome. This suggests we have assembled a highly complete genome with relatively low levels of redundancy. We then performed three rounds of genome annotation by training the annotation program using MAKER2. In total, we annotated 15,225 genes. Among which, 13,435 genes have reciprocal best hits in *D. suzukii*. We used this gene set for downstream comparative genomic and transcriptomic analyses. This high-quality genome makes it possible to study the genomic differences and genome evolution of *D. suzukii* and *D. subpulchrella*, and how they compare to other closely related species.

The genome and assembly of *D. subpulchrella* is now available at NCBI through GCF_014743375.2 or JACOEE01. In total, 15,028 protein-coding genes were annotated, which is very similar to our MAKER2 annotation. 29,037 transcripts were annotated with a median length of 1.9kb, which is similar to the median transcript length of *D. melanogaster*. The median number of transcripts per gene was 1.67, and the median number of exons per transcript was 6.16. In addition, a total of 2481 non-coding genes were annotated. We found that 23.95% of the genome was repetitive sequences using RepeatMasker, which is considerably higher than other *Drosophila* species such as *D. melanogaster* (Kaminker et al., 2002), *simulans* species complex (Chakraborty et al., 2020), and *D. hydei* (Zhao and Begun, 2017). Repetitive regions were comprised of 14.09% retroelements, 0.41% DNA transposons, and 3.01% simple repeats, among other types of repetitive sequences. Notably, 6.82% belongs to the Gypsy family, and 1.33% belongs to R1/LOA/Jockey retrotransposons; it would therefore be an interesting future direction to investigate if the telomere elongation mechanism of *D. subpulchrella* is similar to *D. biarmipes* (Saint-Leandre et al., 2019).

### Adaptive evolution of sensory receptors implicated in *D. suzukii* oviposition site preference

Our behavioral analyses show that changes along the *D. suzukii* lineage, in both chemosensory and mechanosensory systems, are involved in the stepwise transition to ripe fruit preference in *D. suzukii*. Therefore, to investigate the genetic changes that lead to *D. suzukii*’s unique oviposition behavior, we analyzed the repertoire of *Drosophila* olfactory, gustatory, and mechanosensory receptor genes for signals of positive selection using population level genomic data from over 200 *D. suzukii* females. Specifically, we performed a McDonald-Kreitman test (MK) to identify candidate odorant receptors (OR), gustatory receptors (GR), ionotropic receptors (IR), and mechanosensory receptors (MR) evolving under positive selection in *D. suzukii* as compared to *D. subpulchrella, D. biarmipes*, and *D. melanogaster* separately (Figure 3). The MK test infers the presence of positive selection by comparing the numbers of fixed and segregating, synonymous and nonsynonymous substitutions in the genomes of a focal population (McDonald and Kreitman, 1991). While there is previous knowledge on ORs evolving under positive selection in *D. suzukii* as compared to *D. melanogaster* (Ramasamy et al., 2016), our analysis has the additional power of population level data, the additional pairwise comparisons of *D. suzukii* to both *D. biarmipes* and *D. subpulchrella*, and the addition of GRs, IRs, and MRs.

**Figure 3.**
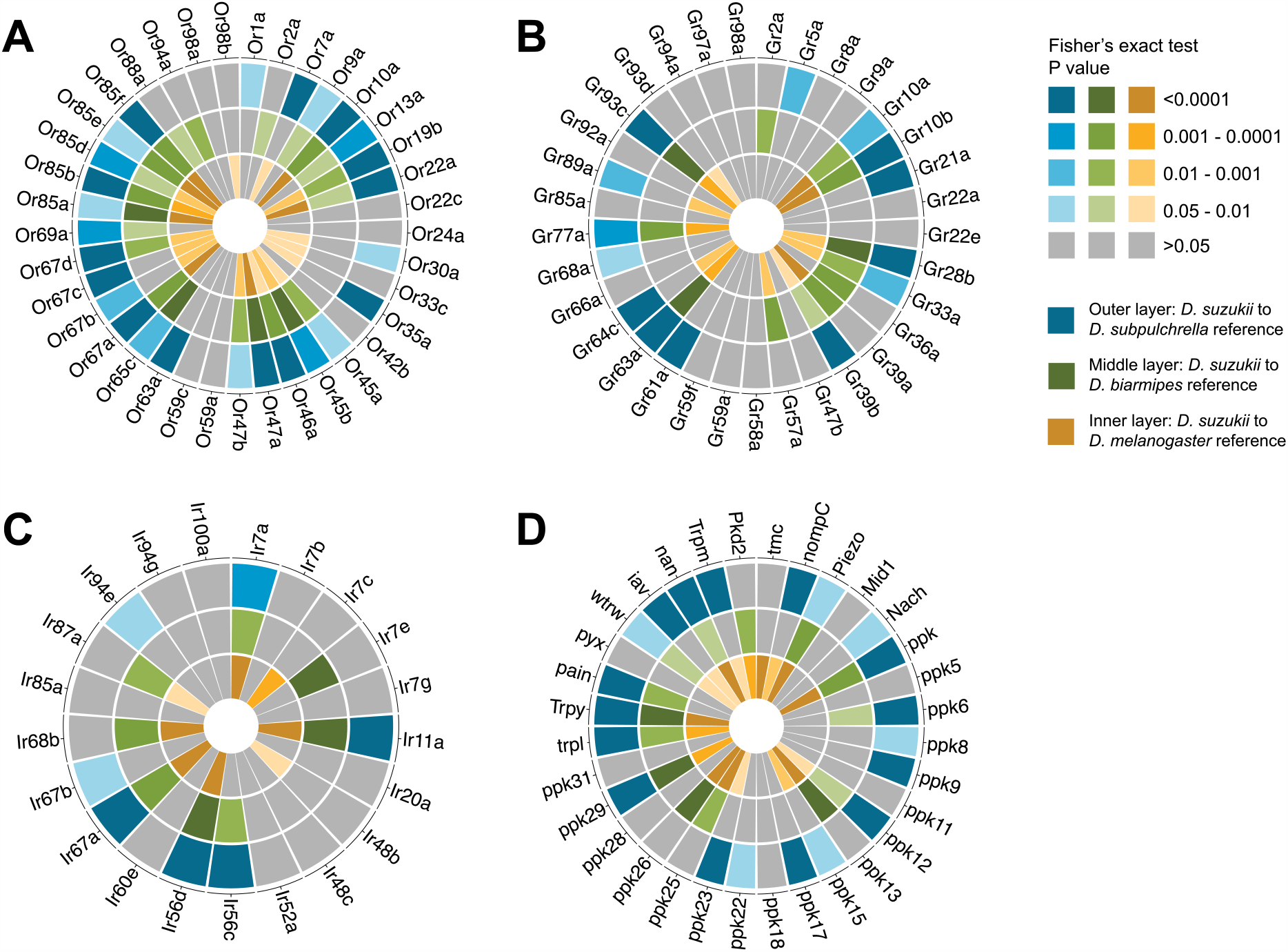
McDonald-Kreitman test for sensory receptors evolving under positive selection in *D. suzukii*, as compared to *D. subpulchrella, D. biarmipes*, and *D. melanogaster*. Odorant receptors (A), gustatory receptors (B), divergent ionotropic receptors (C), and mechanosensory receptors (D) undergoing adaptive evolution in *D. suzukii*. MK test conducted using population data from >200 *D. suzukii* genomes. P-values calculated using Fisher’s exact test.

We used three independent methods to perform the MK test (see methods), and consistently identified about 3000 genes evolving under positive selection in *D. suzukii*, as compared to the other three focal species. This number is very large compared to positively selected genes in other *Drosophila* species when inferred through whole genome and population level genomic analyses, which range from 500-1000 positively selected genes (Begun et al., 2007; Langley et al., 2012; Zhao and Begun, 2017). Specifically, we found that *d*_N_/*d*_S_ is significantly larger than *p*_*N*_*/p*_*S*_, and that both *d*_N_ and *d*_S_ are large (Figure S3). Since *D. suzukii* recently invaded North America, its evolutionary history suggests that the over-representation of positively selected genes may be caused by the bottleneck of invasion and the subsequent increase in population size and likely effective population size (Eyre-Walker, 2002; McDonald and Kreitman, 1991). Even if the number of genes under positive selection is overestimated, one would expect that there is no bias in which gene categories are enriched for signals of selection. For the majority of species pairwise comparisons between the four focal *Drosophila* species, the sensory receptors analyzed are evolving under positive selection in *D. suzukii* to a greater degree than other regions of the genome (Figure 3, Figure S4). This overrepresentation for signals of positive selection in olfactory, gustatory, and mechanosensory receptors suggest that these genes are at least partly contributing to adaptive evolution in *D. suzukii*. On the other hand, there is also a likelihood of false negative results, and sensory genes that are not significant in our MK test may still be important for adaptive evolution, as changes to a small number of amino acids can cause strong shifts in receptor affinity and specificity, a signal that would not be captured by our analysis. Thus, we focus on identifying candidate sensory genes that show expression divergence or adaptive gene evolution, rather than analyzing them as a whole.

Receptors that are under positive selection in *D. suzukii* in each of the three pairwise species comparisons may underlie *D. suzukii*’s novel oviposition behavior (Figure 3). Such genes include *Or22a*, which is involved in host choice evolution in various *Drosophilid* species (Auer et al., 2020; Goldman-Huertas et al., 2015; Mansourian et al., 2018), *Or85a*, a gene implicated in *D. sechellia*’s specialization on morinda fruit (Auer et al., 2020) that is also thought to either be a pseudogene (Hickner et al., 2016) or have changed function in *D. suzukii* (Ramasamy et al., 2016), and *Or85d*, which is linked to the detection of yeast volatiles in *D. melanogaster* (Tichy et al., 2008). Gustatory and ionotropic receptors of interest include *Gr63a*, which is involved in the detection of CO_2_, a volatile emitted by ripening fruit (Jones et al., 2007; Krause Pham and Ray, 2015; Kwon et al., 2007), and *Ir7a*, an ionotropic receptor linked to acetic acid consumption avoidance in *D. melanogaster* (Rimal et al., 2019). Lastly, mechanosensory receptors of note include *nan*, which is involved in substrate stiffness detection during egg-laying in *D. melanogaster* (Jeong et al., 2016; Zhang et al., 2020), *ppk*, a gene involved in acetic acid attraction during oviposition choice in *D. melanogaster* (Gou et al., 2014), and *Trpγ*, which is also linked to CO_2_ detection (Badsha et al., 2012).

### Differential expression of sensory receptors implicated in *D. suzukii* oviposition site preference

To further understand the molecular changes associated with *D. suzukii*’s oviposition preference, we sequenced full transcriptomes and analyzed gene expression data from female adult heads of *D. suzukii, D. subpulchrella, D. biarmipes* and *D. melanogaster*. We analyzed expression across all genes and determined differential gene expression in *D. suzukii* as compared to the three other focal species. Significantly differentially expressed genes in each species comparison and the GO enrichment for these gene sets can be found in supplementary file 3. To determine the strongest candidate genes, we then focused on the same set of sensory receptors included in our MK analysis (Table S6), and built upon our positive selection population genomic analysis to curate a final list of genes which exhibit both significant MK tests (in all three species pairwise comparisons), and a significant difference in gene expression (in at least one species comparison). These criteria generated 15 candidate receptor genes potentially underlying *D. suzukii*’s oviposition behavior (Figure 4). These genes include previously mentioned receptors *Or85a, Or85d, Gr63a*, and *Trpγ*, strengthening the potential role of CO_2_ detection and host fruit specializing in *D. suzukii*’s ripe fruit preference. This list also includes *Ir56d*, another receptor implicated in the response to carbonation and CO_2_ (Sánchez-Alcañiz et al., 2018). Other ORs implicated are *Or10a* and *Or85f*, which are both involved in response to benzaldehydes in *D. melanogaster* (Rollmann et al., 2010), a major volatile emitted by fruits during ripening (Girard and Kopp, 1998). Lastly, candidate MRs include *wtrw*, which is involved in humidity sensation in *D. melanogaster* (Liu et al., 2007), *trpl*, a cation channel shown to mediate gradual dietary shifts in *D. melanogaster* (Zhang et al., 2013), and *Piezo*, which interacts with the receptor *nan*, mentioned above, to sense substrate stiffness differences in *D. melanogaster* (Zhang et al., 2020).

**Figure 4.**
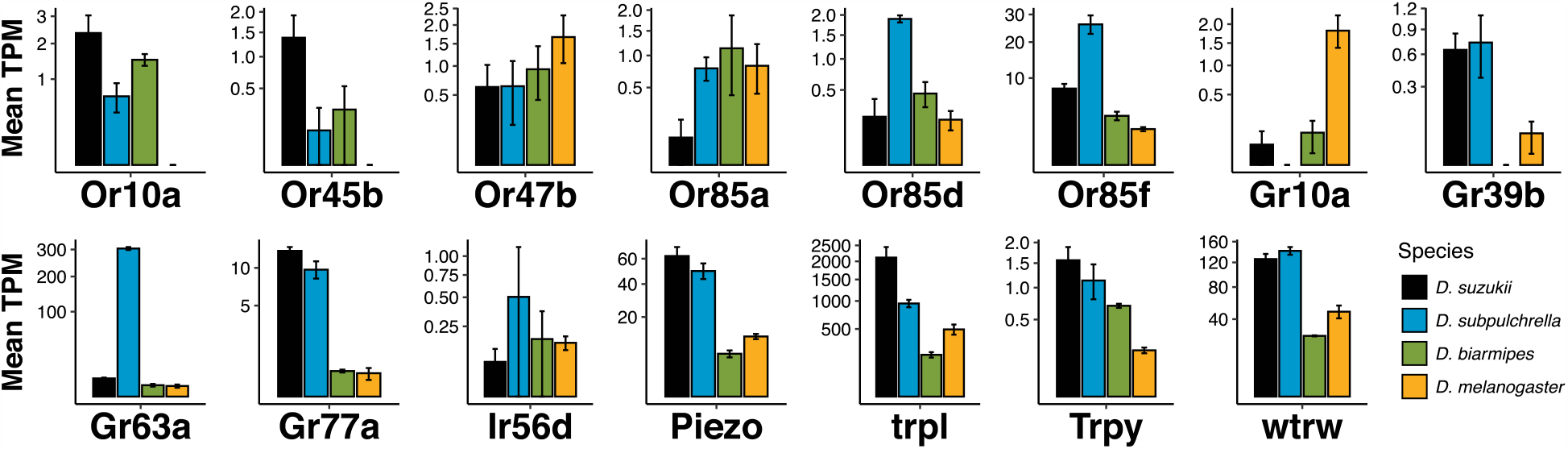
Gene expression of candidate sensory receptors potentially underlying *D. suzukii*’s novel oviposition behavior. Fifteen candidate ORs, GRs, IRs, and MRs were identified as both evolving under positive selection in D. *suzukii* compared to *D. subpulchrella, D. biarmipes*, and *D. melanogaster* and significantly differentially expressed (P < 0.05) in at least one species as compared to *D. suzukii* in the gene expression analysis. The y-axes are mean transcripts per million (TPM) normalized between all species and each y-axis is square root transformed. P-values listed in supplementary tables S2-5 and calculated using species pairwise *t*-tests. Histogram bars are mean ± SD.

It is important to note that our results represent data from a single, heterogeneous tissue, and cannot capture many elements of gene expression evolution along the *D. suzukii* lineage. To address this, we analyzed gene expression in the same set of sensory receptors included in our MK analysis (Table S6) using previously published transcriptome data from the abdominal tip of our four focal species (Crava et al., 2020). Overall, fewer sensory receptor genes were expressed in the abdominal tip (Table S7). However, several of our candidate genes, including *Piezo, nan, wtrw, trpl, Or85f*, and *Gr63a* were expressed at different levels among the four focal species, pointing to potentially interesting future directions. Additionally, to ascertain how well our head RNA-seq dataset captures expression information from relevant sensory organs, we analyzed the correlation between our *D. suzukii* head RNA-seq data and published RNA-seq datasets from the proboscis + maxillary palp and antennae of *D. suzukii* females (Paris et al., 2020) among genes in our sensory receptor gene set (Table S6). We found that the relative expression of sensory genes in our *D. suzukii* head RNA-seq dataset and in the proboscis + maxillary and antennae palp datasets were significantly correlated (Spearman’s r = 0.32 and 0.67, respectively, P = 0.003 and 5.5×10^−12^, respectively) (Figure S5), suggesting that our sample collection method can relatively accurately measure the expression of relevant sensory genes in the context of this study.

While we focus on behavioral and genomic changes to the peripheral sensory system here, we acknowledge that differences in oviposition choice between species could be due to changes in central-brain processing (Seeholzer et al., 2018), and this would be an interesting direction for future studies on *D. suzukii* oviposition preference. In the long-term, it is important to understand how stiffness sensation influences species-specific behavioral differences. We hope the mechanosensory genes observed in this work, such as *Piezo* and *nan*, among other genes (Zhang et al., 2020), will help shed light on the evolution of mechanosensation in this species group.

### CONCLUSIONS

Here we present an investigation of the behavioral patterns, sensory modalities, genetic factors and evolutionary forces contributing to the emergence of ripe fruit preference in *D. suzukii*, a newly invasive and rapidly spreading fruit pest. We show that *D. suzukii* differs from closely related species *D. subpulchrella, D. biarmipes* and *D. melanogaster* in discrete and important ways as it pertains to oviposition preference for whole fruit, and common substances associated with fruit maturation and rotting. As compared to the other *Drosophila* species studied, *D. suzukii* prefers to oviposit in ripe fruit, displays an indifference toward sucrose, an aversion to acetic acid, and a preference for stiff oviposition substrates. The species-specific egg-laying behavior of *D. suzukii* has been shown to be associated with the enlargement and strengthening of the ovipositor allowing females to puncture stiff fruit, a trait shared by its sister species, *D. subpulchrella* (Atallah et al., 2014). Previous work established a stepwise model for the evolution of *D. suzukii* as a pest species in relation to *D. melanogaster* and *D. biarmipes*, focusing on ovipositor size and fruit stage preference (Karageorgi et al., 2017). Our work clarifies and builds upon this model with the addition of empirical fruit preference data in *D. subpulchrella*, a species representing an intermediate step toward the exploitation of ripe fruit as an egg-laying substrate, as well as an in-depth analysis of sensory cues used for the discrimination of fruit maturation stages (Figure 5).

**Figure 5.**
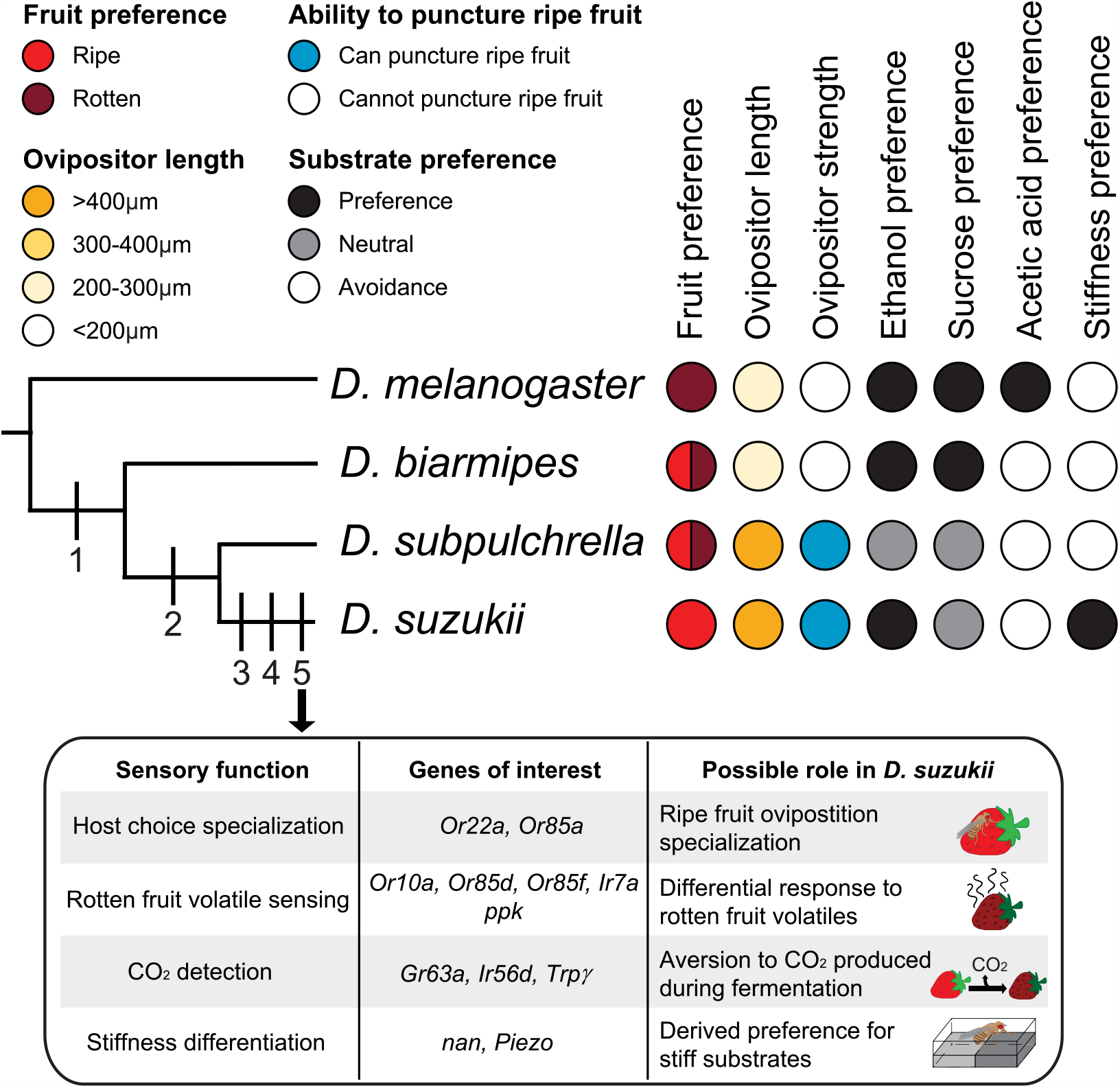
Model of best behavior evolution in *D. suzukii*. The traits acquired step-wise leading to the specialization on ripe fruit in this model are: 1. Relaxation of rotten fruit preference, 2. Enlarged ovipositor and ability to puncture ripe fruits, 3. Ripe fruit specialization, 4. Preference for stiff substrates, and 5. *D. suzukii*-specific molecular sensory changes (highlighted in the inset table). Ovipositor strength and length data from Atallah et al. 2014.

In addition to clarifying the egg-laying preference of *D. subpulchrella*, we generated a high-quality genome for this species. This high-quality genome, together with the genomes of *D. suzukii* (Chiu et al., 2013; Paris et al., 2020) including a PacBio-based assembly (Paris et al., 2020), will benefit evolutionary studies of the *suzukii* species group. Future comparative studies focused on *D. subpulchrella* may help reveal how *D. suzukii* evolved into an invasive pest, while other closely related species did not. Finally, we analyzed the genomic changes associated with *D. suzukii*’s sensory evolution and generated a list of candidate olfactory, gustatory and mechanosensory receptors with signals of both differential gene expression and positive selection (Figure 5). A substantial number of these genes seem to be evolving under positive selection and are differentially expressed in *D. suzukii* as compared to the other species analyzed, despite being recently diverged.

Similar instances of rapid sensory receptor gene evolution can be seen across a wide variety of taxa during the adaptation to new ecological niches. Within the family *Drosophilidae*, sensory receptors have undergone lineage- and species-specific genomic changes driven by positive selection that are linked to ecological specialization (Guo and Kim, 2007). As an example, *Or22a* has undergone several independent molecular evolution events, including instances of neuronal expansion in the morinda fruit specialist, *D. sechellia* (Auer et al., 2020; Dekker et al., 2006) and the *Pandanus* fruit specialist, *D. erecta* (Linz et al., 2013), and instances of gene deletion in the herbivorous fly, *S. flava* (Goldman-Huertas et al., 2015). Additionally, *Or22a* displays high levels of genetic differentiation among ancient populations of *D. melanogaster* specializing on marula fruit (Mansourian et al., 2018). The implication of rapid sensory receptor evolution in the adaptation to new ecological niches is pervasive beyond the family *Drosophilidae*, such as in the unique pheromone system of orchid bees (Brand et al., 2015), the specialization of human blood feeding in *Aedes aegypti* mosquitos (McBride et al., 2014), and ecotype adaptation across phylogenetically and ecologically diverse mammals (Hayden et al., 2010). This representation across the Kingdom Animalia highlights the importance of sensory receptor gene evolution in mediating ecological shifts.

Previous work on our identified candidate genes in *D. melanogaster* and other *Drosophilid* species suggests that changes in CO_2_ detection, stiffness differentiation, rotten fruit volatile sensing, and host choice specialization in part underlie *D. suzukii*’s oviposition preference for ripe fruit (Figure 5). Further work is required to truly understand the functional implications of the genetic changes seen in *D. suzukii*, and the results outlined here are a valuable resource for future studies aimed at understanding the behavioral and molecular basis of pest activity in this species.

## METHODS

### Fly Stocks and husbandry

All flies were reared on standard cornmeal medium at 24°C, 55% relative humidity, on a 12-hour light-dark cycle (lights on at 8:00am). Egg-laying experiments were conducted under the same conditions. For behavioral assays we used a set a wild type strains: Canton S for *D. melanogaster*, the genome strain raj3* (Chen et al., 2014) for *D. biarmipes*, #NGN6 from Japan for *D. subpulchrella* (Ehime Stock Center), and the genome strain WT3 (WT3, Chiu et al., 2013) for *D. suzukii*. We tested different lines within species and found all the behaviors are consistent. For example, in *D. melanogaster, w* ^*1118*^ and Canton S show the same results, suggesting that the traits tested here are likely to be fixed in each species. For genomic analyses we used the genome reference strain of *D. melanogaster* (BDSC #2057, Adams et al., 2000), the genome strain of *D. biarmipes* (raj3*) (Chen et al., 2014), the genome strain of *D. suzukii* (WT3, Chiu et al., 2013), and our lab inbred strain of the wild caught line *D. subpulchrella* (Ehime Stock Center #NGN6).

### Egg-laying assays

#### Whole fruit 2-choice oviposition assay

Flies were collected as virgins and aged for 7-8 days in food vials containing approximately 10 males and 10 females. For each trial 10 females and 5 males were placed in a mesh experimental cage (10 × 10 × 10 inch, BugDorm 4F2222) which contained both a whole ripe and whole rotten strawberry without anesthesia using an aspirator. Flies were allowed to lay eggs for 19 hours from late afternoon to the next morning, after which the total number of eggs laid in each fruit was counted, and an oviposition preference index (OPI) was calculated as follows: (number of eggs on ripe fruit – number of eggs on rotten fruit)/(number of eggs on ripe fruit + number of eggs on rotten fruit). Ripe strawberries (always of the same variety) were purchased from a local supermarket the day of the experiment, and rotten strawberries (same variety, purchased from the same supermarket) were allowed to rot in a 24°C, 55% relative humidity room for four days prior to the experiment. Only intact fruits without any damage were used in experiments. Between 10 and 12 replicate assays were performed for each species. In total, 45 assays were performed. Cut-ripe fruit vs. rotten fruit assays (Figure S1) were performed in the same way as whole-fruit choice assays, except the ripe fruit was cut in half before the trail, and the exposed flesh was placed facing up in the behavioral chamber.

#### Substrate 2-choice oviposition assays

For all substrate choice assays flies were collected as virgins, separated by sex, and aged separately for 3-4 days in food vials. 2-3 days prior to the experiment, males and females were placed in a new food vial supplemented with yeast paste (1.5ml ddH_2_O + 1g live active yeast) to mate and produce eggs. For each trial 3-4 females were placed in a custom laser cut egg-laying chamber (see description below) containing an agarose pad of the experimental substrate, and an agarose pad of the control substrate. Flies were inserted through a trap door with an aspirator without the use of anesthesia. Flies were allowed to lay eggs for 19 hours, after which the total number of eggs laid on each agarose pad was counted, and the OPI was calculated as follows: (number of eggs on experimental substrate – number of eggs on control substrate)/(number of eggs on experimental substrate + number of eggs on control substrate). Only assays where flies had laid a total of 10 eggs were included in the final analyses, and between 15 and 32 replicate assays were performed for each species at each concentration point. In total, 905 assays were performed.

##### Chamber design

Egg-laying chambers for substrate choice assays were custom made using laser cut acrylic plastic. Each chamber contained two separate egg-laying arenas separated by 6 mm of plastic, so two trials could be conducted in one chamber. Each arena contained two wells, into which agarose substrates were poured. Wells measured 32×45×12.7 mm and were divided by 5 mm of plastic. Each arena contained a trap door through which flies could be inserted without the use of anesthesia. A 100×75 mm glass sheet covers the entire chamber to allow light into the chamber, and to prevent flies from escaping.

##### Sucrose

For sucrose preference assays, experimental substrates were 1% agarose and contained 1% ethanol and increasing concentrations (200mM, 400mM or 600mM) of sucrose (ThermoFisher #S5-3); control substrates were 1% agarose and contained 1% ethanol.

##### Ethanol

For ethanol preference assays, experimental substrates were 1% agarose and contained 75mM sucrose and increasing concentrations (3%, 5%, and 7%) of ethanol (ThermoFisher #BP2818); control substrates were 1% agarose and contained 75mM sucrose.

##### Acetic Acid

For acetic acid preference assays, experimental substrates were 1% agarose and contained 75mM sucrose, 1% ethanol and increasing concentrations (1.5%, 2.5%, and 3.5%) of acetic acid (ThermoFisher #A465-250); control substrates were 1% agarose and contained 75mM sucrose and 1% ethanol.

##### Agarose

For agarose/stiffness preference assays, experimental substrates contained 75mM sucrose, 1% ethanol and increasing concentrations (1%, 1.5%, and 2%) of agarose (Lonza SeaKem LE Agarose #50001); control substrates were 0.25% agarose and contained 75mM sucrose and 1% ethanol.

#### Statistics

All statistical analyses were performed using R (RStudio version 1.2.1335). For the whole fruit oviposition assays, a Wilcoxon signed rank test was used with the null hypothesis set to 0, signifying no preference. For the substrate gradient preference oviposition assays, a linear regression approach was performed for each substrate using the lme4 package in R to find the overall preference difference between species across the concentrations tested. To determine if the preference curve across the three concentrations differed significantly between the four focal species, we used the lm function in the lme4 package with the response term of OPI and predictor terms of the cross of concentration and species. The reference group, the species to which the other species are compared, was manipulated using the relevel function in base R to perform pairwise comparisons between the preference curves of each species. Effectively, this sets each of the four different species as the baseline OPI response to the concentration tested and compares this baseline to the other species analyzed. P-values refer to the difference between pairwise comparisons of the y-intercepts of each species’ preference curve across the three concentrations. The P-values represent the level of significance for the difference between the slope of each species’ regression and 0. Error bars in all figures are mean ± standard error of the mean (SEM) unless otherwise noted.

### Genomic analysis

#### D. subpulchrella genome library preparation and sequencing

##### D. subpulchrella inbred line generation

To generate an inbred line for PacBio sequencing, *D. subpulchrella* flies (Ehime Stock Center #NGN6) were inbred via sib-mating for ten generations to generate the strain denominated “*D. subpulchrella* 33 F10 #4”.

##### DNA extraction and Sequencing

We extracted DNA from adult females following the protocol of Chakraborty et al. (Chakraborty et al., 2016). The DNA was sheared using 20 plunges of 21-gauge needle and size selected using the 30kb lower cutoff size on Blue Pippin size selection system (Sage Science). 30kb SMRTbell template library was prepared from the size selected DNA and was sequenced on 4 SMRTcells of Pacific Biosciences Sequel I platform. We also sequenced this sheared DNA sample with 150 bp paired-end library on Illumina Hiseq 4000. All sequencing was performed at UCI GHTF.

##### Genome assembly

We generated 45.3 Gb of long reads, in which 50% of the sequence is contained within sequences longer than 33.8kb (the longest sequence is 160 kb) and 149.40 million 150 bp paired-end Illumina reads. The reads were corrected and assembled with canu v1.7 using genomeSize=220M (Koren et al., 2017). The assembly was polished twice with arrow (smrtanalysis v5.1.0) using the long reads and twice with Pilon using the Illumina reads (Walker et al., 2014). The size of the final assembly was 265 Mb, 50% of which is contained within reads that are 11.59 Mb or longer (assembly contig N50 11.59 Mb). The genome assembly can be found on NCBI through WGS project# JACOEE01 or BioProject# PRJNA557422. We ran RepeatMasker against HMM-Dfam_3.2 lib to estimate the number of repetitive sequences in *D. subpulchrella*.

#### D. subpulchrella genome annotation

To evaluate the genome assembly quality, we used BUSCO (Simão et al., 2015) to estimate the proportion of the 2799 Diptera orthologous genes and the 303 eukaryotic genes that were completely or partially assembled in the genome. We then used the MAKER2 (Holt et al., 2011) genome annotation pipeline for genome annotation. To improve the annotation quality, we trained the HMM for three times before using it for the final annotation (Zhao and Begun, 2017) After that, we used OrthoMCL (Li et al., 2003) to find homologous genes between *D. subpulchrella* and the other *Drosophila* species. For multiple-copy genes, we assigned their orthologous genes by using reciprocal best hits through BLASTP. The NCBI annotation can be found through GenBank assembly accession: GCA_014743375.2 and RefSeq assembly accession: GCF_014743375.2.

#### McDonald-Kreitman test for positive selection

A high-quality population genome dataset was aligned to the *D. suzukii* genome using bowtie2. After that we called bi-allelic SNPs using ANGSD version 0.920 (Korneliussen et al., 2014). On average the coverage of locations with SNPs is 178. We used bi-alleles that met the following criteria: minimum mapping quality of 30, minimum allele frequency of 0.05, and a minimum coverage larger than 10. We then created alternative references using the set of SNPs. We re-extracted the coding sequence of each gene from alternate references, then re-aligned using PRANK with the –codon function for each *D. suzukii* gene and their orthologous gene in *D. melanogaster, D. biarmipes*, and *D. subpulchrella*. We only carried out MK tests for genes that showed at least one variant in each of four categories, polymorphic, fixed, synonymous, and nonsynonymous (Begun et al., 2007). For genes that passed the above criteria we carried out unpolarized McDonald–Kreitman tests using the MK.pl script (Begun et al., 2007), and a version of our own python scripts independently, followed by a Fisher’s exact test for each gene within each species pairwise comparison. A third method of MK test using SNPs directly from individuals was also used for confirming the results. For each gene, we estimated the proportion of adaptive amino acid fixations (α) (Smith and Eyre-Walker, 2002) and the Direction of Selection (DoS) index (Stoletzki and Eyre-Walker, 2011).

Lists of ORs, GRs, IRs and MRs were compiled using FlyBase assigned gene groups for *D. melanogaster* and their orthologous genes in the other three species were inferred using OrthoMCL or BLASTP. The ORs, GRs, IRs, and MRs with orthologs in each of the three species, and that met the MK test criteria described above were used for genomic and transcriptomic analyses (Table S6). Short protein sequences do not fit the MK test criteria due to lack of polymorphisms and are therefore not included in our downstream analyses.

Simulations were used to compare the average degree of adaptive evolution occurring in receptor gene groups to the genome-wide average. We randomly sampled gene sets of identical size to the receptor gene set of interest for each separate species comparison (*D. suzukii – D. melanogaster, D. suzukii – D. biarmipes*, and *D. suzukii – D. subpulchrella*) and computed the mean Fisher’s exact test P-value for that random gene set. Simulations were run 10,000 times for each set of interest, and P-values represent the proportion of the simulated distribution of means below the observed mean P-value for the receptor gene set of interest.

#### RNA library preparation, and sequencing

We generated head transcriptomes from female *D. suzukii, D. melanogaster, D. biarmipes*, and *D. subpulchrella*. All individuals were mated and precisely 3-days-old. Female flies were very briefly anesthetized with CO_2_ and heads were collected with a clean razor blade. Dissections were performed within a 1-hour window always at the same circadian zeitgeber time (ZT1-ZT2; with light turning on at 8 am, and the dissections being performed between 9 am and 10 pm). We collected 3 biological replicates for each species, each sample contained 15 heads. Dissected heads were immediately transferred into a low retention Eppendorf tube containing 100 μL TRIzol (Invitrogen) and RNA was extracted immediately post dissection.

All RNA extractions were performed according to TRIzol manufacturer protocol and immediately followed by a DNase treatment using the TURBO DNase from Invitrogen. RNA quality was assessed by a Bioanalyzer run of an Agilent Eukaryote Total RNA Pico chip while RNA quantity was measured with a Nanodrop One (ABI). About 1 μg total RNA was used for library preparation. Libraries were fragmented and enriched for mRNA using NEBNext Poly(A)^+^ Magnetic Isolation Module (NEB #E7490) and prepared using NEBNext Ultra II Directional RNA Library Prep Kit (NEB #E7765) and single indexing from the NEBNext Multiplex Oligos kit (NEB #E7555L) following manufacturer protocol including beads size selection for 200 bp inserts. Library quality was first assessed on Agilent D1000 ScreenTapes for Tapestation and then by Qubit and Agilent Bioanalyzer. Finally, 150 bp paired-ends libraries were sequenced on an Illumina Nextseq500 platform.

#### Identification of differentially expressed sensory receptor genes

Adaptors and low-quality bases from RNA-seq reads were trimmed using Trimmomatic (Bolger et al., 2014) using the setting LEADING:1 TRAILING: 1 SLIDINGWINDOW:20:25 and MINLEN:36. Bowtie2 (Langmead and Salzberg, 2012) was then used to align reads to the reference genome of the species being analyzed. Gene and transcript expression levels (TPM: Transcripts per million) were then quantified using StringTie (Pertea et al., 2015). We then obtained a list of homologous genes of *D. suzukii, D. biarmipes*, and *D. subpulchrella* using OrthoMCL and reciprocal best hits. To compare gene expression between replicates from different species we used TMM (trimmed mean M-values) normalized TPM values. TMM normalization was implemented with the R package edgeR (Robinson et al., 2010). We then conducted a Log_2_ transformation across all replicates and calculated a P-value using a Student’s *t*-test between *D. suzukii* and each of the other focal species to determine if each gene is differential expressed between two species. We specifically queried the expression patterns of sensory receptor genes inferred by homology with *D. melanogaster* sensory receptor genes extracted from FlyBase, using the same gene list as that described for the MK test above (Table S6) (Thurmond et al., 2019). GO enrichment of significantly differentially expressed genes (P < 0.01) was performed using PANTHER (Mi et al., 2019).

#### Evaluation of sensory receptor gene expression in other relevant tissues

Raw RNA-seq reads from the female abdominal tip of the four focal species were downloaded from the Genbank SRA database (BioProject# PRJNA526247) (Crava et al., 2020). Raw reads of adult female proboscis + maxillary palp and female antennae for *D. suzukii* were downloaded from NCBI SRA SRR10716767 and SRR10716770, respectively (Paris et al., 2020). Reads were aligned and analyzed similarly to that described above, with the only difference that no statistical analysis was performed because the data do not have biological replicates.

Correlation between the head RNA-seq dataset and the maxillary palp + proboscis and antennae RNA-seq datasets was done by first ranking the expression of sensory receptor genes (Table S6) in each of the three datasets. Correlation of the rankings was calculated using the Spearman correlation coefficient in R. Genes that are not expressed were ranked as 1. We used rankings to compare these datasets because the relative raw expression levels could be influenced by technical differences that do not reflect biologically meaningful information.

## DATA AVAILABILITY

All single-molecule sequence data and RNA-seq data have been deposited to the NCBI Sequence Read Archive (SRA) and can be found under BioProject accession PRJNA557422. The *D. subpulchrella* genome assembly has been deposited in the NCBI Assembly database under accession JACOEE01. The vcf file for the population genome data can be found in figshare: https://doi.org/10.6084/m9.figshare.c.5237618. The genomes and SNPs used in the MK analysis, and raw behavioral data can be found at the GitHub page https://github.com/LiZhaoLab/D.suzukii_genomes_and_analysis.

## ACKNOWLEDGEMENTS

The authors are grateful for the help from many scientists: Vikram Vijayan and Gaby Maimon for help with design and construction of chambers for egg-laying assays used in Figure 2 and Figure S2, and for helpful comments and suggestions; Precision Instrumentation Technologies (PIT) at Rockefeller for help with construction of behavioral chambers; Samuel Khodursky for suggestions on the gene expression analysis; Clara Drew for help with the statistical analyses; the Ehime Stock Center and Dr. Masayoshi Watada from Ehime University for the help with fly stocks; and members of the Zhao lab for helpful discussions. We thank Terence Murphy and Françoise Thibaud-Nissen from NIH Eukaryotic Genome Annotation team for their help with annotation and making the data readily available for the general community.

## AUTHOR CONTRIBUTION

S.M.D, L.Z., and N.S. conceived the study. S.M.D performed the behavioral analyses with the help of M.K. C.B.L and AG performed cut-ripe fruit behavioral experiments for the revision of the manuscript. S.M.D generated the genome strain of *D. subpulchrella* and performed RNA sequencing library construction. M.C. and J.J.E. sequenced the *D. subpulchrella* genome. A.A., K.M.L., and J.C.C. generated the *D. suzukii* population genome data. L.Z., S.M.D., A.G., J.P., and N.S. performed the computational analysis. S.M.D. and L.Z. wrote the manuscript with the input from all authors.

## SUPPLEMENTARY FIGURES & TABLES FOR

**Figure S1.**
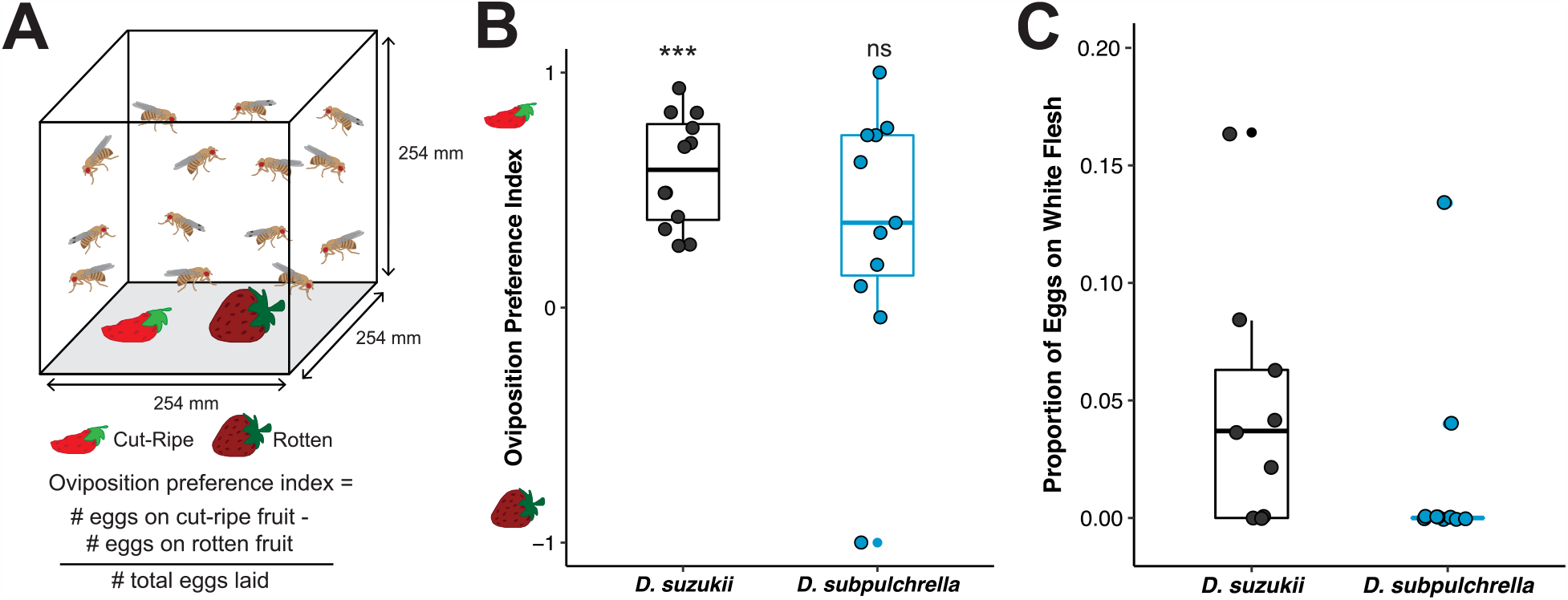
Cut-ripe vs. rotten fruit preference in *D. suzukii* and *D. subpulchrella*. A. Schematic of two-choice oviposition preference assay. 10 females and 5 males were placed in a cage with a cut-ripe (with flesh facing up) and rotten strawberry for 19 hours. Number of eggs on each fruit was then counted. B. Oviposition preference for cut-ripe fruit (positive OPI) vs. rotten strawberry (negative OPI) in *D. suzukii* and *D. subpulchrella*. C. Proportion of eggs laid on the white, exposed flesh of the cut strawberry in comparison to the intact skin of the ripe strawberry. Each data point represents one experimental trial and data dispersion is represented by a boxplot. Preference P-values were calculated using a Wilcoxon signed rank test against a theoretical value of 0 (no preference). ***, p≤0.001; **, p≤=0.01; *, p≤0.05; ns, p>0.05 for all future figures.

**Figure S2.**
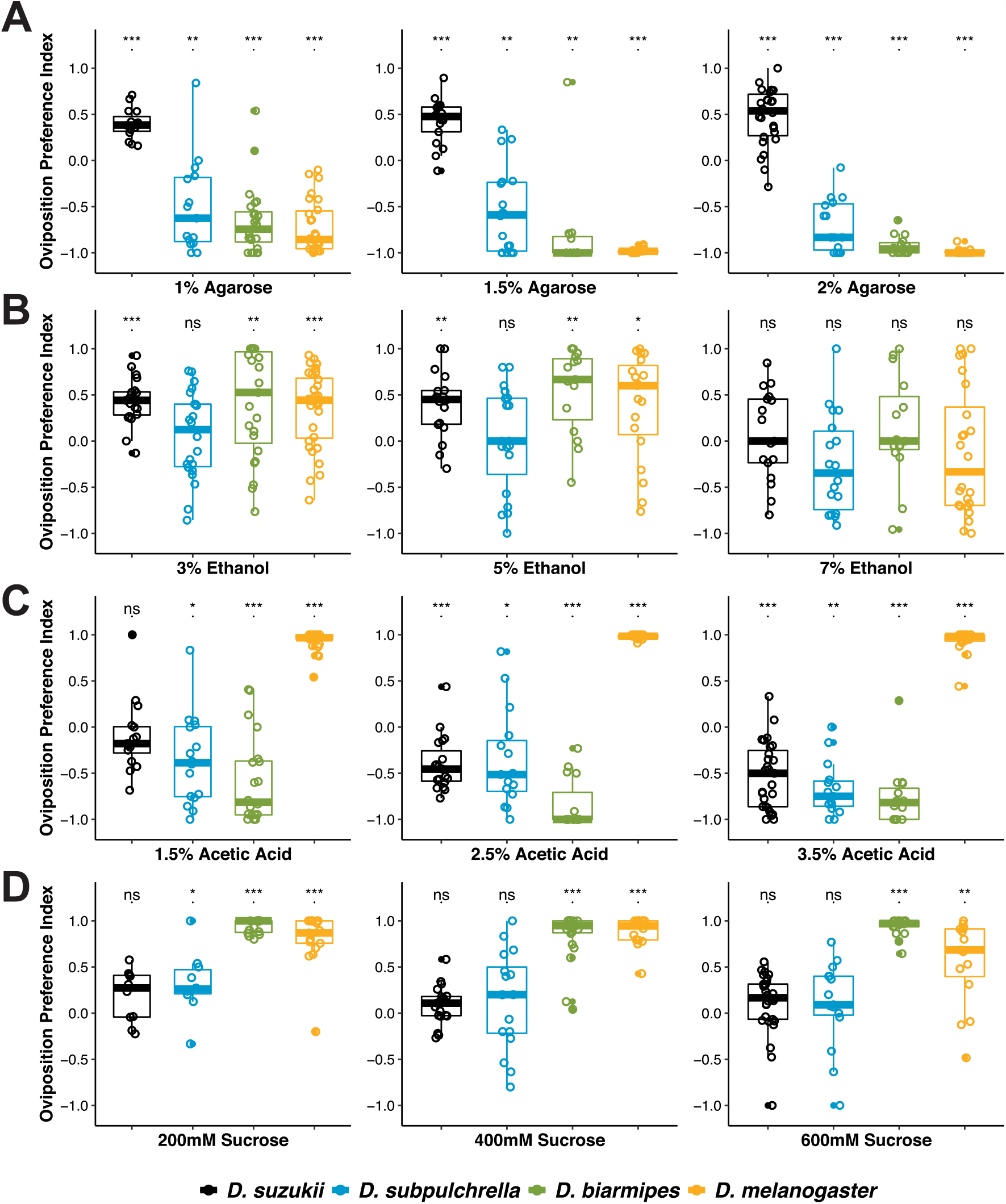
Oviposition preference at each concentration point. Species specific oviposition preference at each separate concentration tested of agarose (A), ethanol (B), acetic acid (C) and sucrose (D). Each data point represents one experimental trial and data dispersion is represented by a boxplot. Preference P-values were calculated using a Wilcoxon signed rank test against a theoretical value of 0 (no preference).

**Figure S3.**
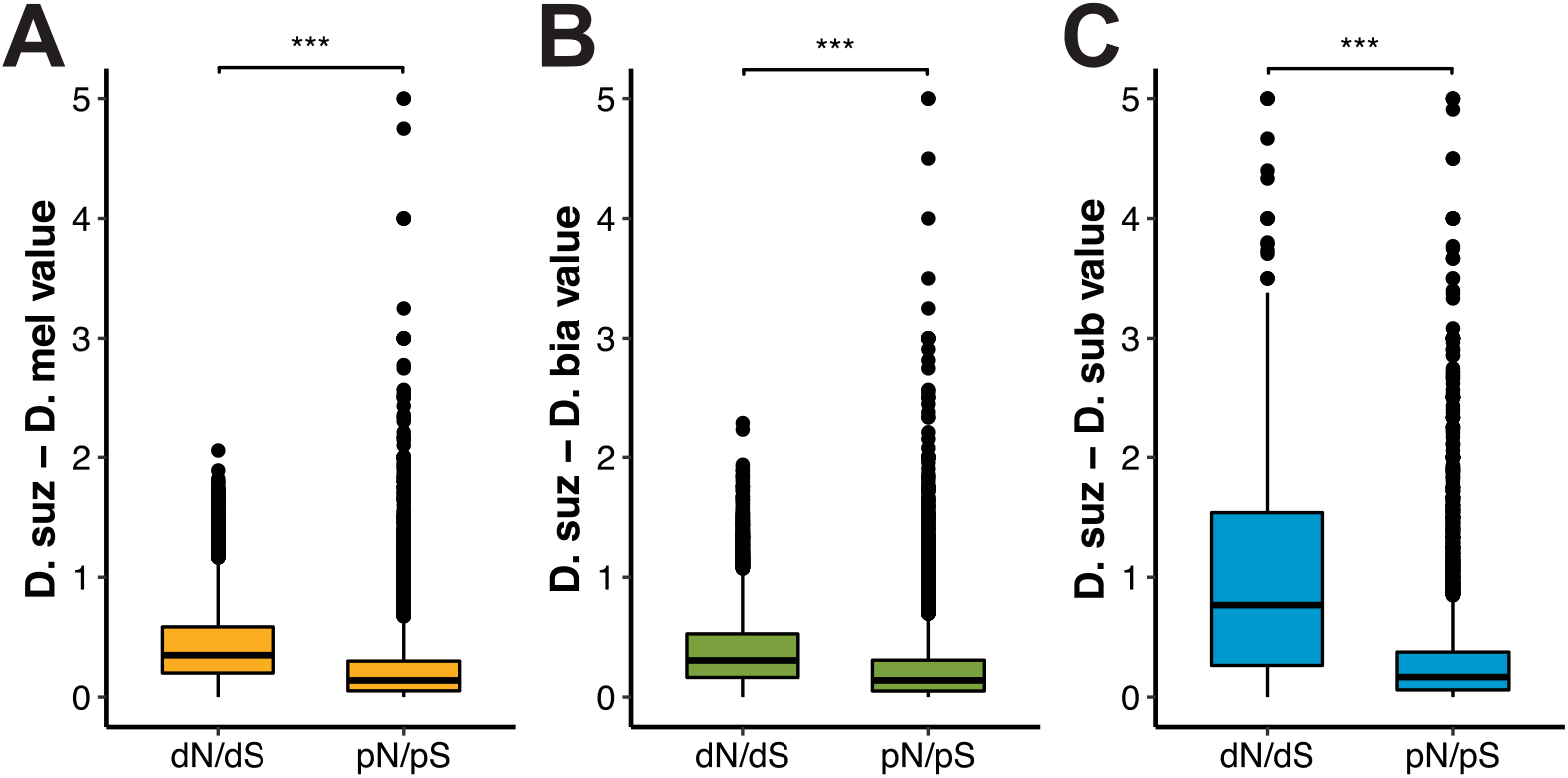
Distribution of *d*_*N*_*/d*_*S*_ and *p*_*N*_*/p*_*S*_ from genome wide McDonald-Kreitman test. *d*_*N*_*/d*_*S*_ is significantly larger than *p*_*N*_*/p*_*S*_ across the genome in all three species pairwise comparisons: *D. suzukii* to *D. melanogaster* (A), *D. suzukii* to *D. biarmipes* (B), and *D. suzukii* to *D. subpulchrella* (C). Y-axes have a cutoff of 5. However, all values were included for statistical analyses and P-values were calculated through pairwise *t*-tests.

**Figure S4.**
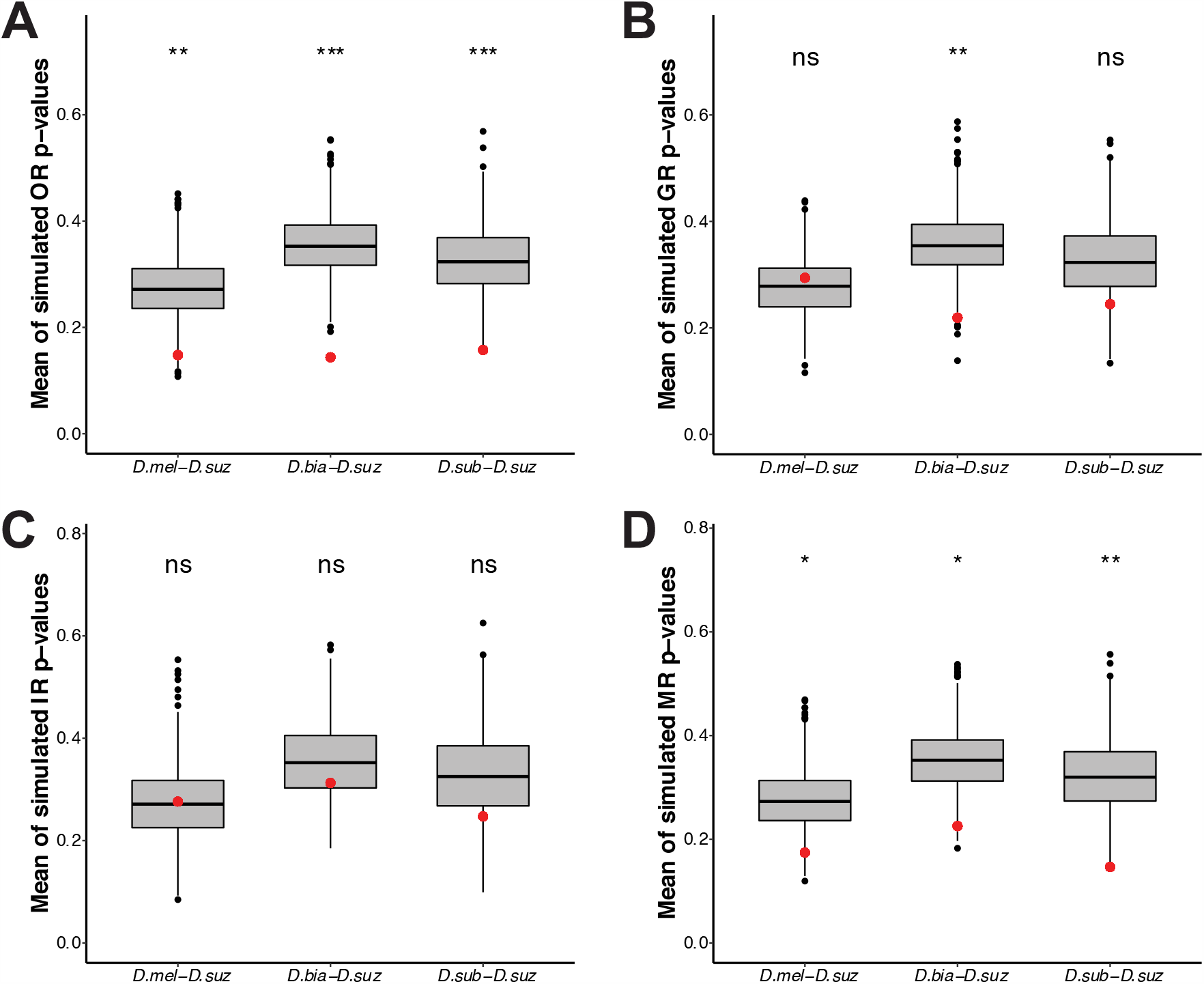
Sensory receptor genes are evolving under positive selection to a greater extent than the genome as a whole. To generate distributions of simulated mean p-values for ORs (A), GRs (B), IRs (C) and MRs (D) for each species comparison, a random group of genes of the same number as the group of sensory receptor gene groups was selected 10,000 times, and the mean of the random gene group was plotted. Red dots represent the actual mean P-value for the sensory receptor gene group within each species comparison. Significance labels represent the proportion of the distribution lower than the actual mean P-value (red dot).

**Figure S5.**
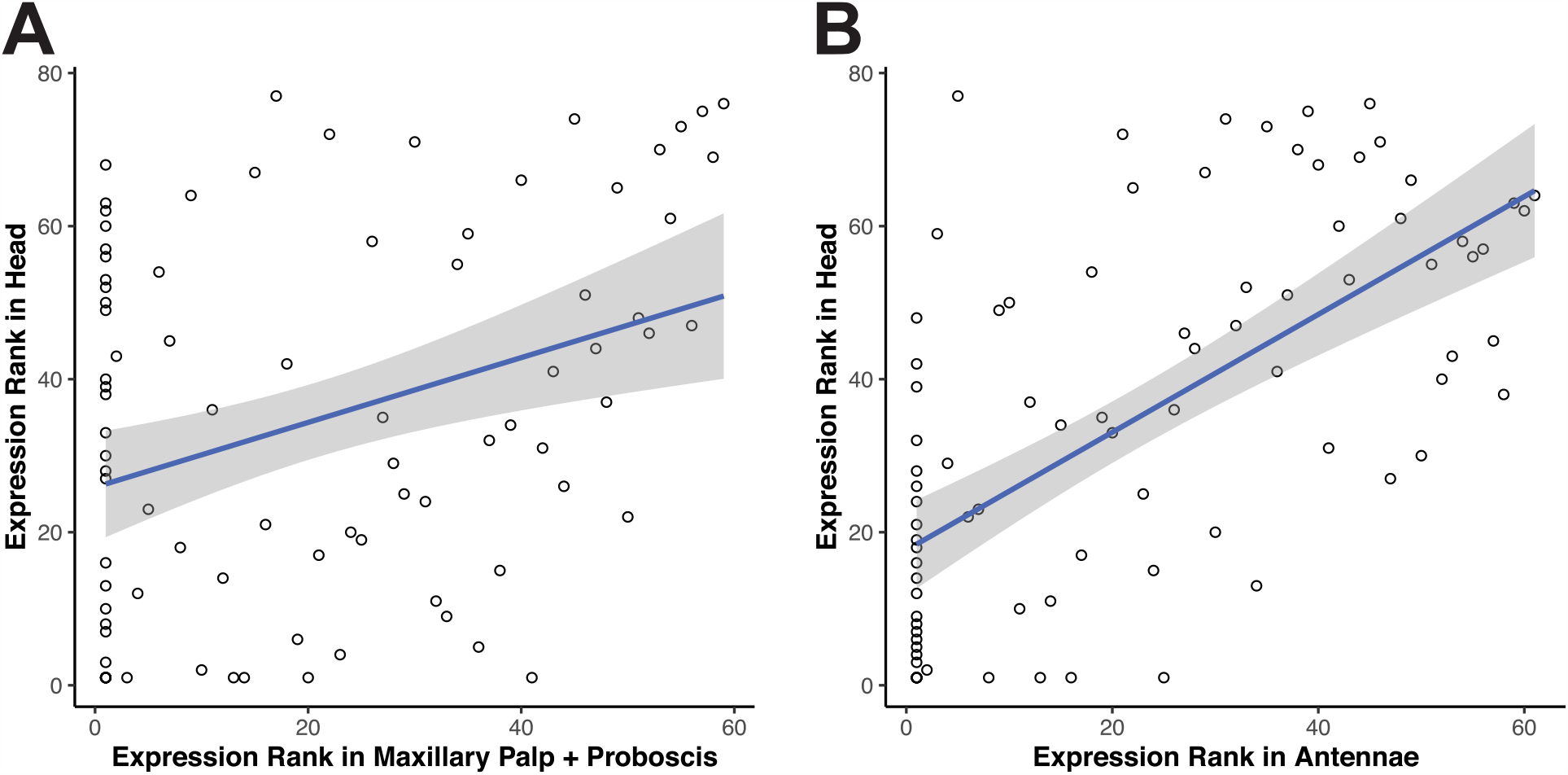
Correlation of expression ranking among female *D. suzukii* head and sensory organs. The relative expression for each gene in our sensory receptor gene set (Table S6) was ranked from lowest to highest in RNA-seq datasets from the maxillary palp + proboscis and antennae and compared to the ranking of expression in our head RNA-seq dataset.

**Table S1.** Properties of the chromosome-level assembly of the *D. subpulchrella* genome. The scaffolds listed in the table have at least 6 genes that are reciprocal best hits of *D. melanogaster* genes. Scaffold names starting with “alt_” denote alternate haplotype contigs.

| *D. subpulchrella*<br>scaffold name | Length (Mb) | *D. melanogaster*<br>chromosome<br>homolog | Best hit gene number<br>with homologous<br>chromosome |
| --- | --- | --- | --- |
| Seq405 | 29.50 | 3L | 1985 |
| Seq361 | 22.17 | 2R | 1807 |
| Seq355 | 20.23 | 3R | 1686 |
| Seq407 | 29.67 | X | 1572 |
| Seq360 | 26.21 | 2L | 1532 |
| Seq354 | 11.59 | 3R | 794 |
| alt_Seq3 | 1.79 | 2L | 106 |
| Seq23 | 2.26 | 4 | 56 |
| alt_Seq10 | 0.75 | 2R | 46 |
| Seq24 | 2.16 | 2R | 41 |
| alt_Seq14 | 0.76 | 2L | 14 |
| alt_Seq417 | 0.47 | 2L | 11 |
| Seq15 | 4.47 | X | 9 |
| alt_Seq9 | 0.08 | 3R | 7 |
| Seq36 | 1.62 | 2L | 6 |

**Table S2.** OR gene differential expression P-values. OR genes that were included in the MK test were analyzed for differential expression in *D. suzukii* as compared to *D. melanogaster, D. biarmipes*, and *D. subpulchrella*, separately. P-values were calculated through pairwise *t*-tests on Log_2_ transformed and normalized TPM between *D. suzukii* and the other species.

| Gene Name | *D. mel* - *D. suz* | *D. bia* - *D. suz* | *D. sub* - *D. suz* |
| --- | --- | --- | --- |
| Or10a | 0.031 | 0.076 | 0.058 |
| Or13a | 0.065 | 0.053 | 0.531 |
| Or19b | 0.282 | 0.632 | 0.662 |
| Or1a | 0.010 | 0.500 | 0.519 |
| Or22a | 0.194 | 0.459 | 0.765 |
| Or22c | 0.506 | 0.080 | 0.310 |
| Or24a | 0.272 | 0.939 | 0.913 |
| Or2a | 0.414 | 0.725 | 0.465 |
| Or30a | 0.506 | 0.106 | 1.000 |
| Or33c | 0.157 | 0.277 | 0.529 |
| Or35a | 0.136 | 0.283 | 0.220 |
| Or42b | 0.196 | 0.632 | 0.338 |
| Or45a | 0.084 | 0.123 | 0.414 |
| Or45b | 0.058 | 0.292 | 0.030 |
| Or46a | 0.069 | 0.155 | 0.335 |
| Or47a | 0.405 | 0.005 | 0.169 |
| Or47b | 0.019 | 0.162 | 0.739 |
| Or59a | 0.355 | 0.141 | 1.000 |
| Or59c | 0.876 | 0.022 | 0.016 |
| Or63a | 0.039 | 0.024 | 0.059 |
| Or65c | 0.031 | 0.070 | 0.026 |
| Or67a | 1.000 | 0.026 | 0.467 |
| Or67b | 0.087 | 0.060 | 0.060 |
| Or67c | 1.000 | 0.681 | 0.241 |
| Or67d | 0.639 | 0.144 | 0.519 |
| Or69a | 0.135 | 0.088 | 0.787 |
| Or7a | 0.543 | 0.792 | 0.385 |
| Or85a | 0.054 | 0.105 | 0.014 |
| Or85b | 0.171 | 0.083 | 0.584 |
| Or85d | 0.288 | 0.039 | 0.007 |
| Or85e | 0.843 | 0.092 | 0.053 |
| Or85f | 0.059 | 0.968 | 0.001 |
| Or88a | 0.042 | 0.042 | 0.529 |
| Or94a | 0.296 | 0.289 | 0.519 |
| Or98a | 0.351 | 0.028 | 0.029 |
| Or98b | 0.782 | 0.926 | 0.285 |
| Or9a | 0.021 | 0.976 | 0.037 |

**Table S3.**
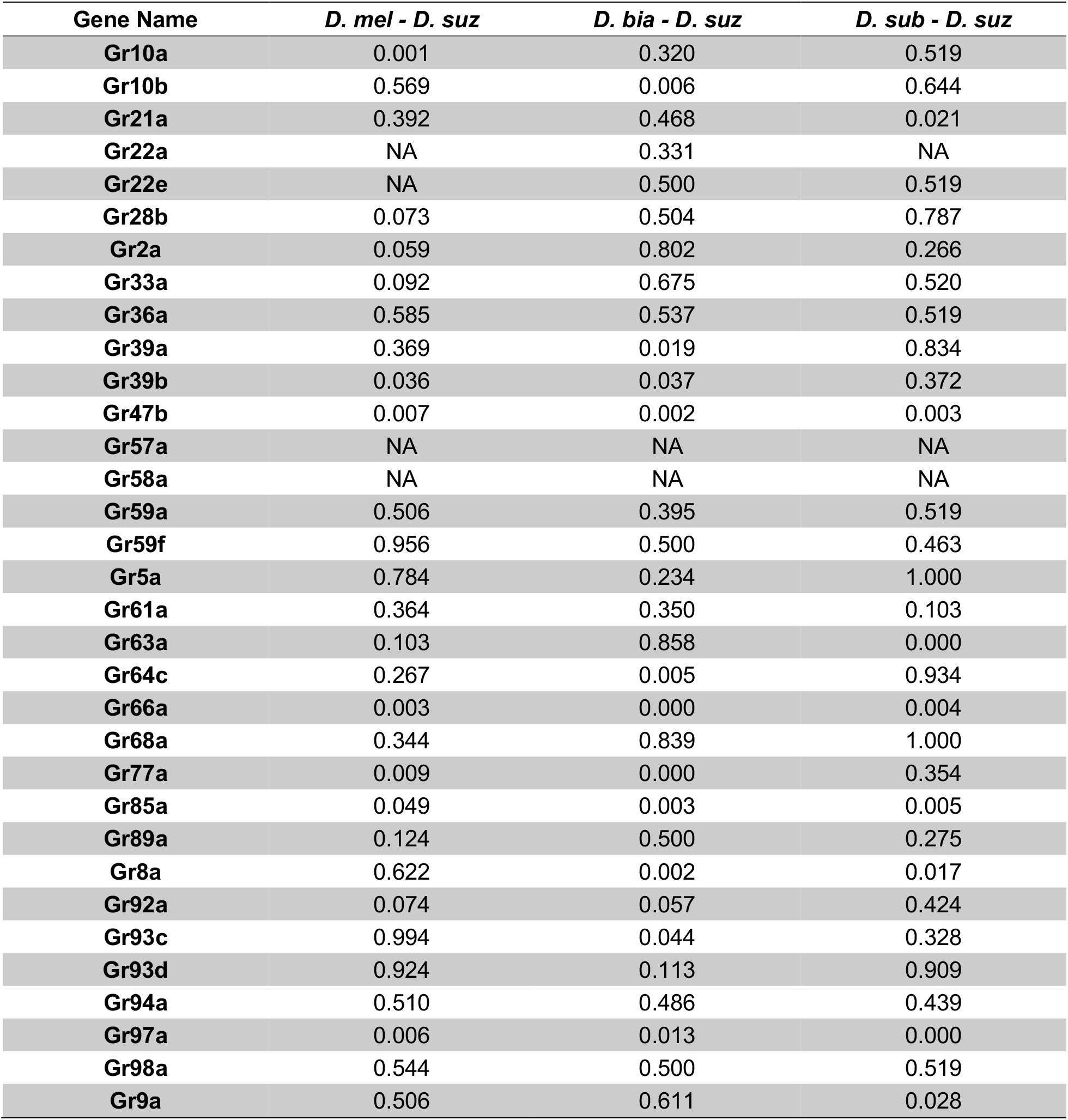
GR gene differential expression P-values. GR genes that were included in the MK test were analyzed for differential expression in *D. suzukii* as compared to *D. melanogaster, D. biarmipes*, and *D. subpulchrella*, separately. P-values were calculated through pairwise *t*-tests on Log_2_ transformed and normalized TPM between *D. suzukii* and the other species. NA – Not Applicable, gene is not expressed in either of the species in the comparison.

**Table S4.**
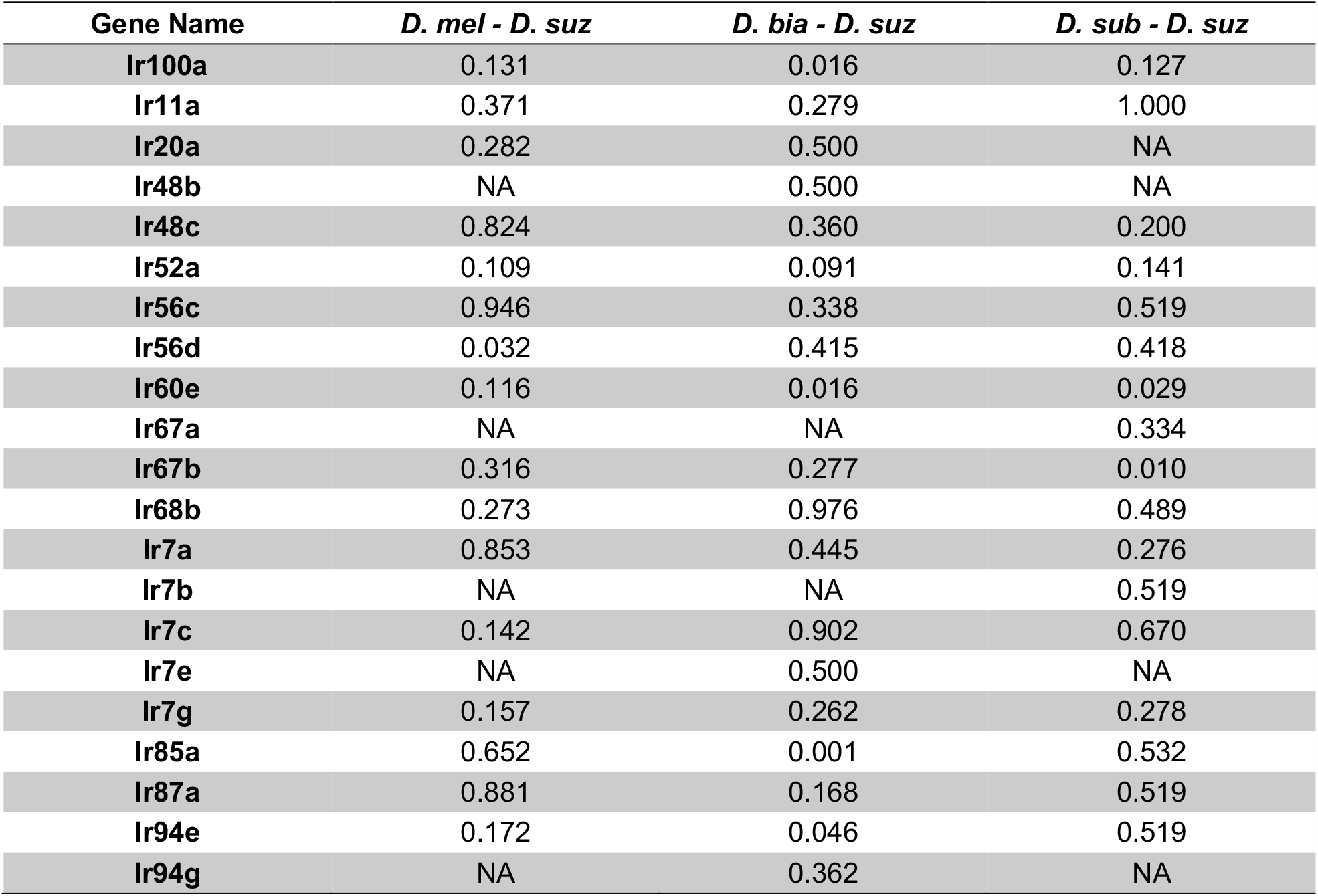
IR gene differential expression P-values. IR genes that were included in the MK test were analyzed for differential expression in *D. suzukii* as compared to *D. melanogaster, D. biarmipes*, and *D. subpulchrella*, separately. P-values were calculated through pairwise *t*-tests on Log_2_ transformed and normalized TPM between *D. suzukii* and the other species. NA – Not Applicable, gene is not expressed in either of the species in the comparison.

| Gene Name | *D. mel* - *D. suz* | *D. bia* - *D. suz* | *D. sub* - *D. suz* |
| --- | --- | --- | --- |
| lr100a | 0.131 | 0.016 | 0.127 |
| lr11a | 0.371 | 0.279 | 1.000 |
| lr20a | 0.282 | 0.500 | NA |
| lr48b | NA | 0.500 | NA |
| lr48c | 0.824 | 0.360 | 0.200 |
| lr52a | 0.109 | 0.091 | 0.141 |
| lr56c | 0.946 | 0.338 | 0.519 |
| lr56d | 0.032 | 0.415 | 0.418 |
| lr60e | 0.116 | 0.016 | 0.029 |
| lr67a | NA | NA | 0.334 |
| lr67b | 0.316 | 0.277 | 0.010 |
| lr68b | 0.273 | 0.976 | 0.489 |
| lr7a | 0.853 | 0.445 | 0.276 |
| lr7b | NA | NA | 0.519 |
| lr7c | 0.142 | 0.902 | 0.670 |
| lr7e | NA | 0.500 | NA |
| lr7g | 0.157 | 0.262 | 0.278 |
| lr85a | 0.652 | 0.001 | 0.532 |
| lr87a | 0.881 | 0.168 | 0.519 |
| lr94e | 0.172 | 0.046 | 0.519 |
| lr94g | NA | 0.362 | NA |

**Table S5.** MR gene differential expression P-values. MR genes that were included in the MK test were analyzed for differential expression in *D. suzukii* as compared to *D. melanogaster, D. biarmipes*, and *D. subpulchrella*, separately. P-values were calculated through pairwise *t*-tests on Log_2_ transformed and normalized TPM between *D. suzukii* and the other species. NA – Not Applicable, gene is not expressed in either of the species in the comparison.

| Gene Name | *D. mel* - *D. suz* | *D. bia* - *D. suz* | *D. sub* - *D. suz* |
| --- | --- | --- | --- |
| iav | 0.008 | 0.002 | 0.060 |
| Mid1 | 0.049 | 0.775 | 0.049 |
| Nach | 0.101 | 0.084 | 0.257 |
| nan | 0.239 | 1.000 | 0.857 |
| nompC | 0.068 | 0.149 | 0.112 |
| pain | 0.001 | 0.036 | 0.581 |
| Piezo | 0.077 | 0.005 | 0.149 |
| Pkd2 | 0.465 | 0.000 | 0.307 |
| ppk | 0.163 | 0.545 | 0.057 |
| ppk11 | 0.328 | 0.751 | 0.456 |
| ppk12 | 0.932 | 0.200 | 0.576 |
| ppk13 | NA | 0.043 | NA |
| ppk15 | 0.084 | 0.003 | 0.350 |
| ppk17 | 0.027 | 0.688 | 0.276 |
| ppk18 | 0.329 | 0.792 | 0.051 |
| ppk22 | 0.637 | 0.001 | 0.255 |
| ppk23 | 0.506 | 0.013 | 0.083 |
| ppk25 | 0.204 | 0.008 | 0.030 |
| ppk26 | 0.506 | 0.717 | 0.953 |
| ppk28 | 0.137 | 0.084 | 0.344 |
| ppk29 | 0.866 | 0.053 | 0.150 |
| ppk31 | 0.011 | 0.133 | 0.024 |
| ppk5 | 1.000 | 0.000 | 0.329 |
| ppk6 | 0.506 | 0.084 | 0.277 |
| ppk8 | 0.591 | 0.946 | 0.195 |
| ppk9 | 0.506 | 0.294 | 0.039 |
| pyx | 0.506 | 0.013 | 0.021 |
| tmc | 0.506 | 0.076 | 0.037 |
| trpl | 0.010 | 0.001 | 0.018 |
| Trpm | 0.078 | 1.000 | 0.343 |
| Trpy | 0.041 | 0.649 | 0.914 |
| wtrw | 0.145 | 0.001 | 0.001 |

**Table S6.**
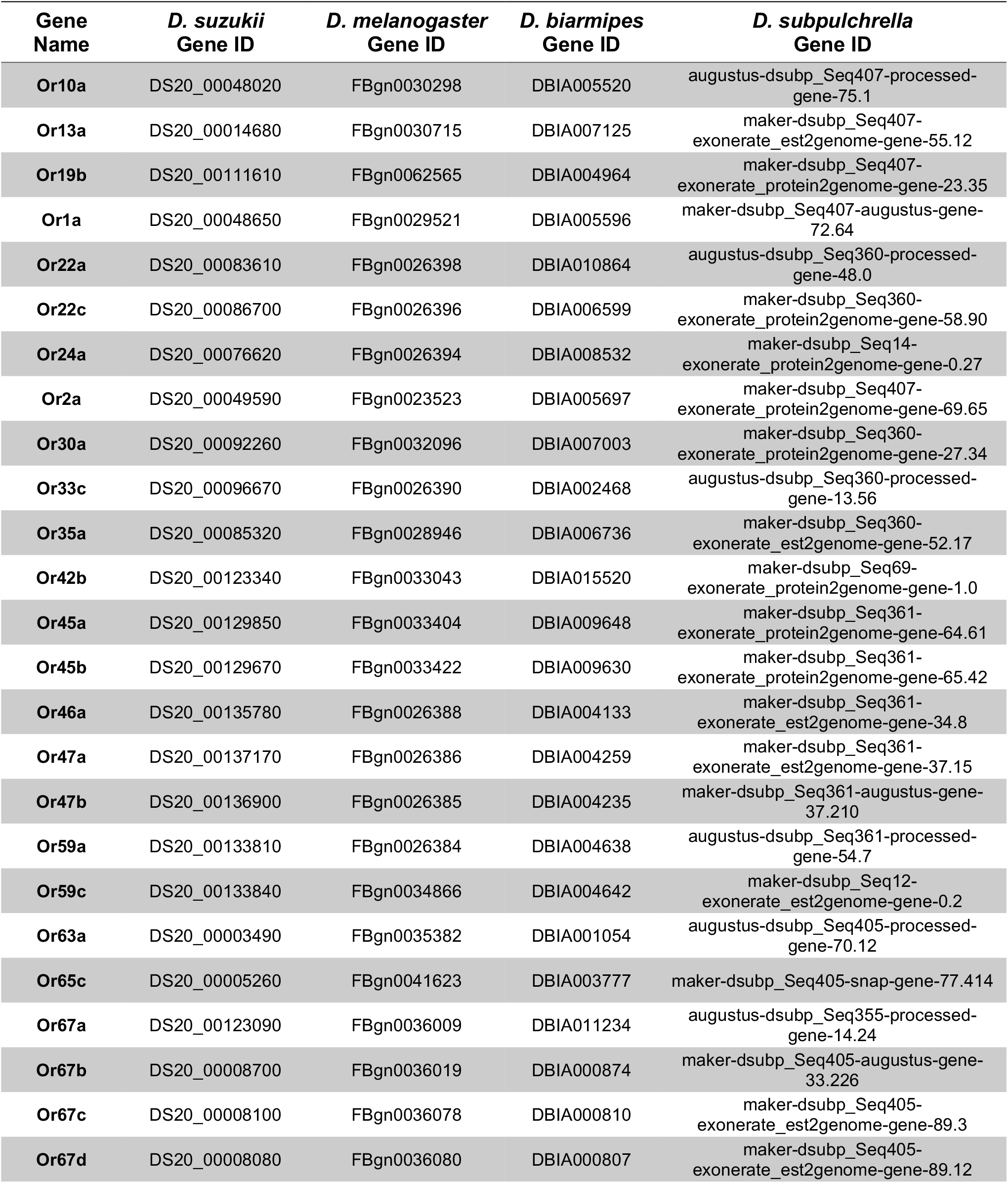
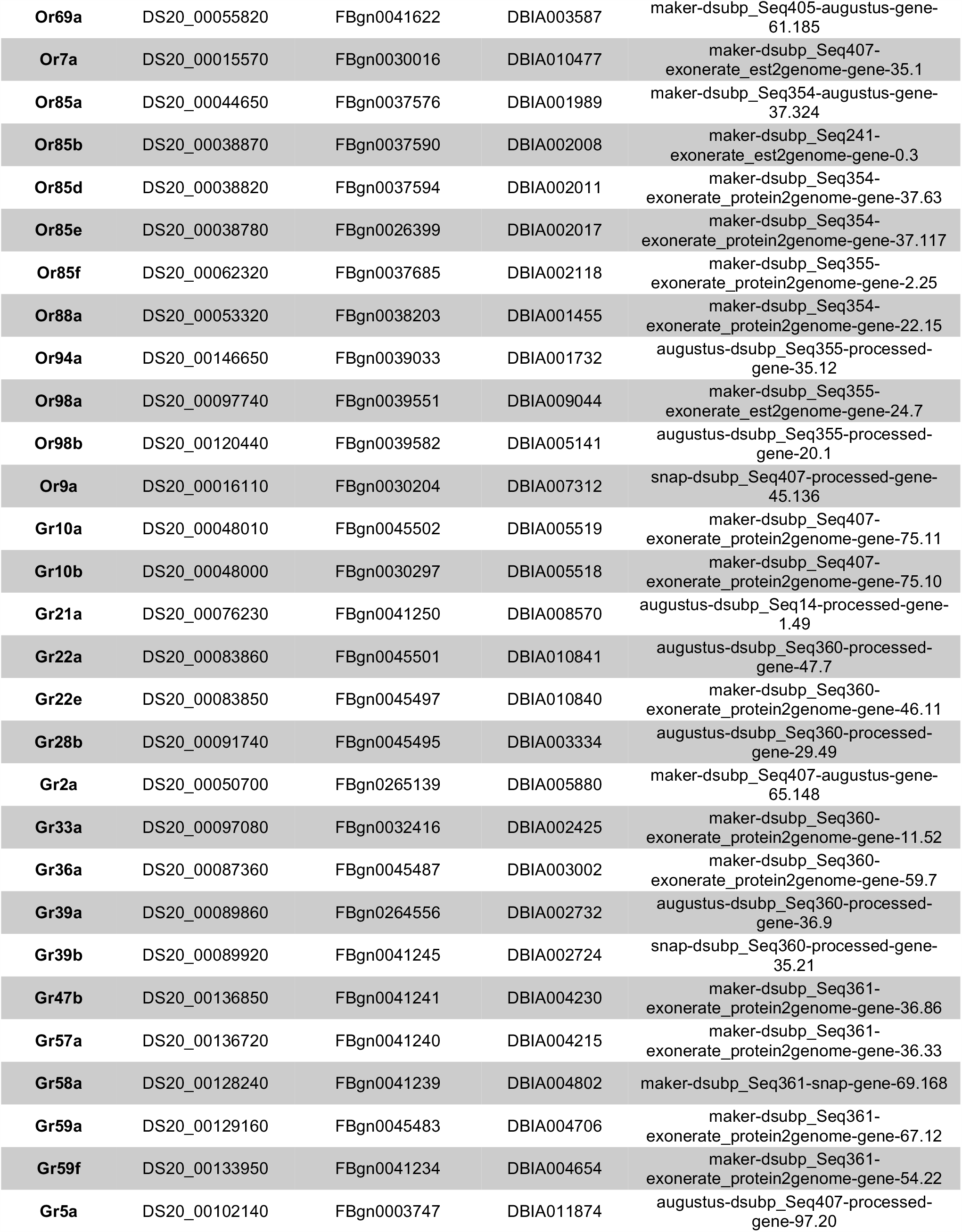
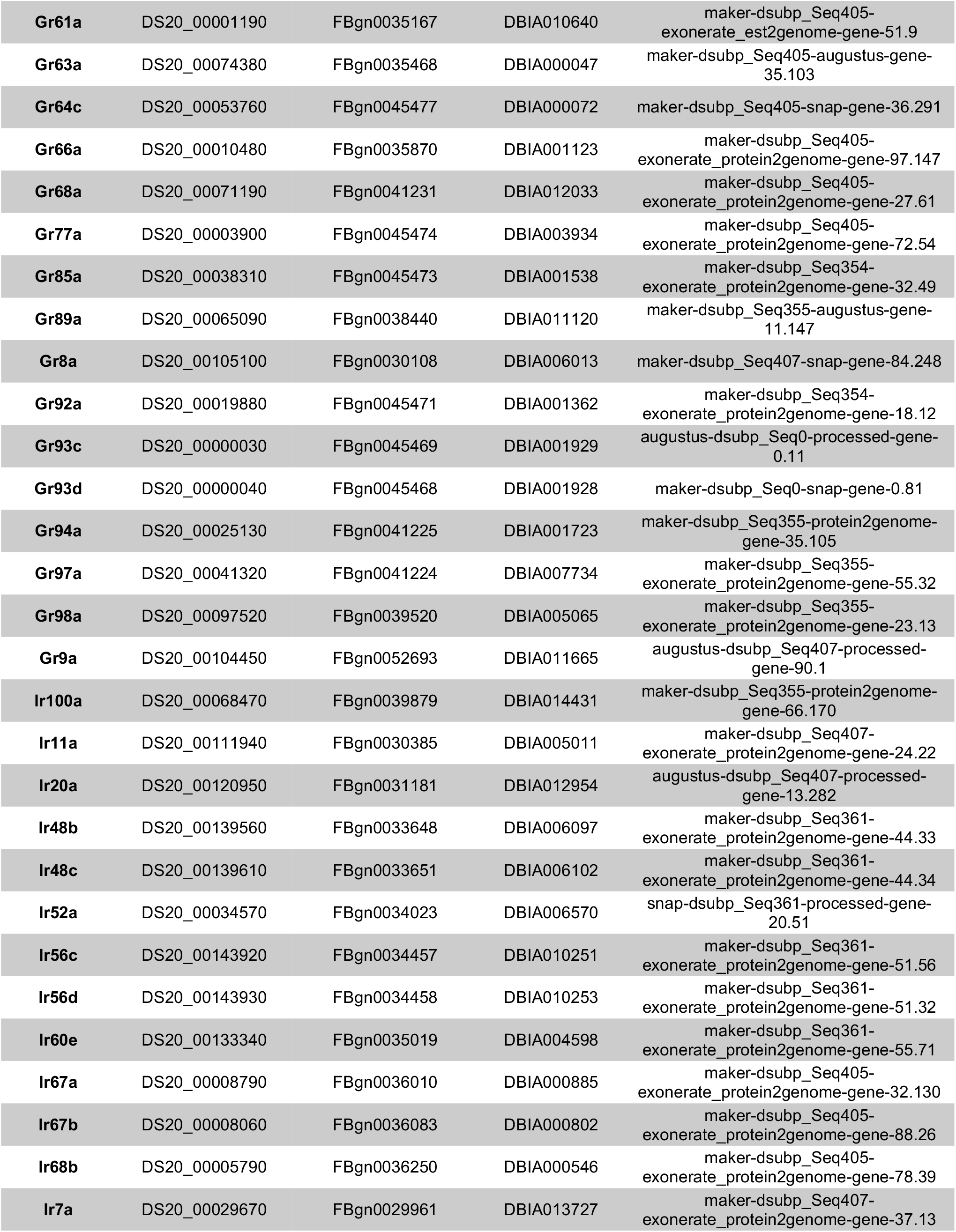
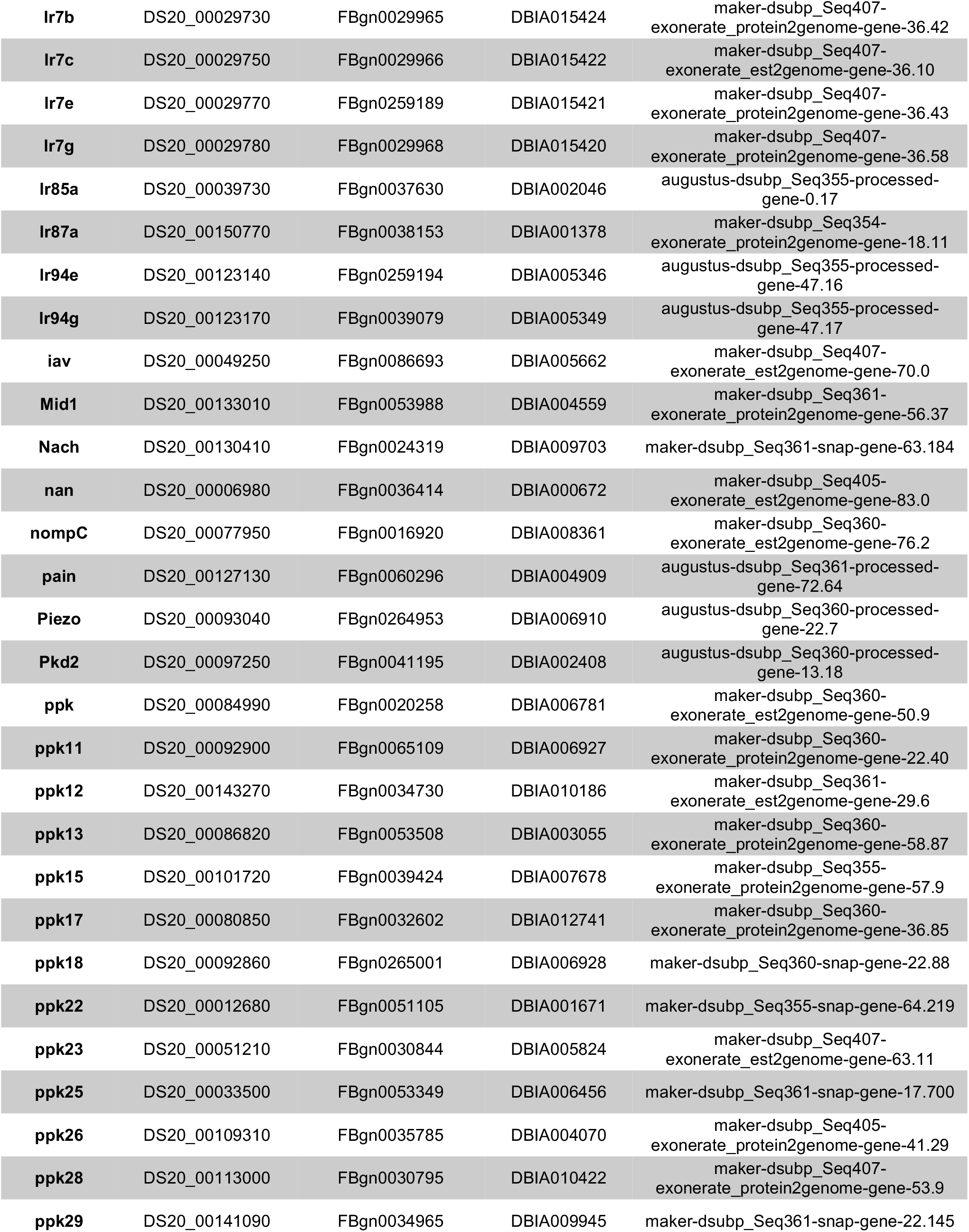
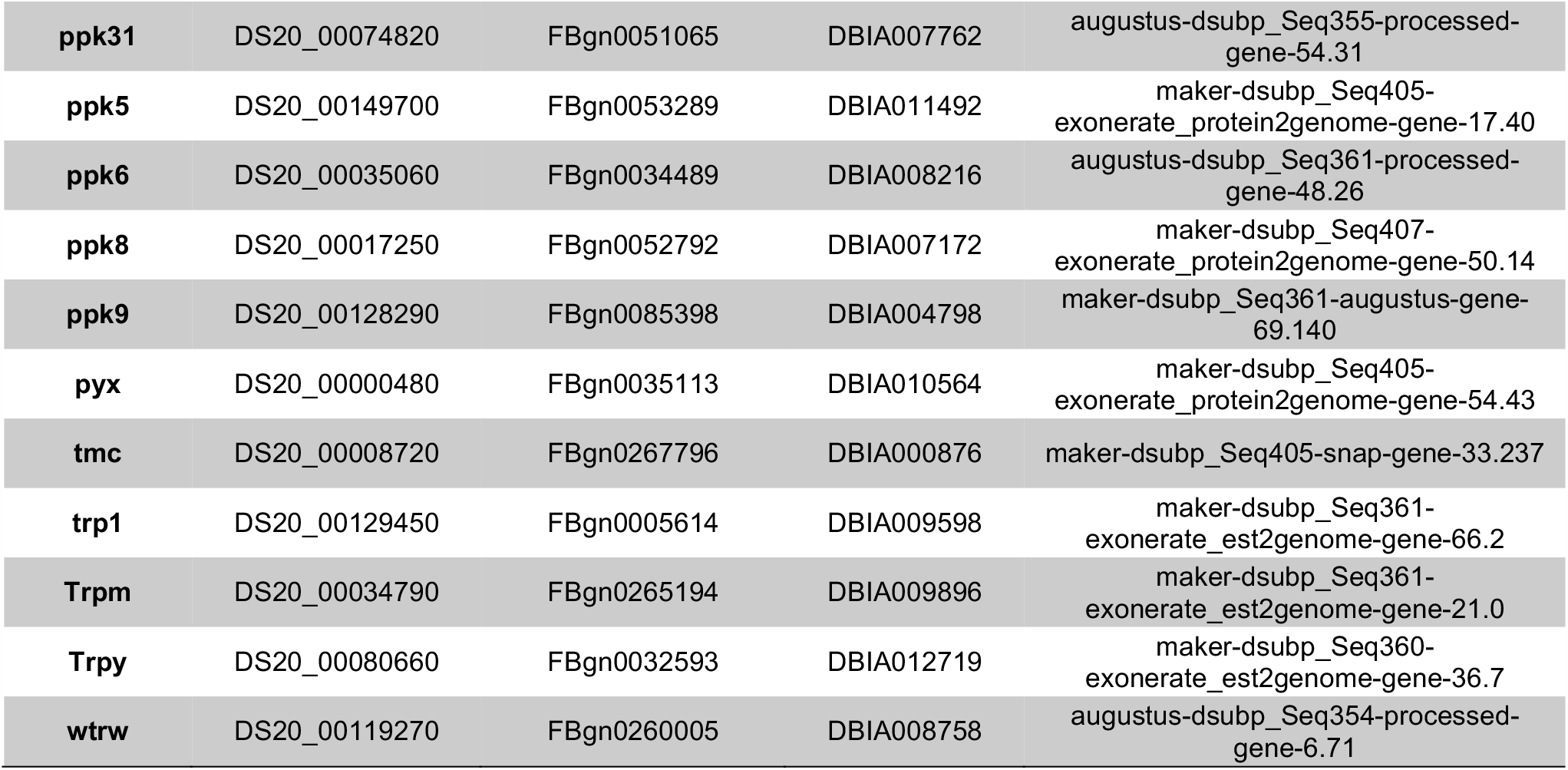
Sensory receptor genes used in genomic and transcriptomic analyses. Ortholog information for OR, GR, IR, and MR genes used for the four focal species. Only genes included in the McDonald-Kreitman (MK) test are listed here. Details of MK test criteria and ortholog matching in methods.

| **Gene Name** | ***D. suzukii***<br>**Gene ID** | ***D. melanogaster***<br>**Gene ID** | ***D. biarmipes***<br>**Gene ID** | ***D. subpulchrella***<br>**Gene ID** |
| --- | --- | --- | --- | --- |
| **Or10a** | DS20_00048020 | FBgn0030298 | DBIA005520 | augustus-dsubp_Seq407-processed-gene-75.1 |
| **Or13a** | DS20_00014680 | FBgn0030715 | DBIA007125 | maker-dsubp_Seq407-exonerate_est2genome-gene-55.12 |
| **Or19b** | DS20_00111610 | FBgn0062565 | DBIA004964 | maker-dsubp_Seq407-exonerate_protein2genome-gene-23.35 |
| **Or1a** | DS20_00048650 | FBgn0029521 | DBIA005596 | maker-dsubp_Seq407-augustus-gene-72.64 |
| **Or22a** | DS20_00083610 | FBgn0026398 | DBIA010864 | augustus-dsubp_Seq360-processed-gene-48.0 |
| **Or22c** | DS20_00086700 | FBgn0026396 | DBIA006599 | maker-dsubp_Seq360-exonerate_protein2genome-gene-58.90 |
| **Or24a** | DS20_00076620 | FBgn0026394 | DBIA008532 | maker-dsubp_Seq14-exonerate_protein2genome-gene-0.27 |
| **Or2a** | DS20_00049590 | FBgn0023523 | DBIA005697 | maker-dsubp_Seq407-exonerate_protein2genome-gene-69.65 |
| **Or30a** | DS20_00092260 | FBgn0032096 | DBIA007003 | maker-dsubp_Seq360-exonerate_protein2genome-gene-27.34 |
| **Or33c** | DS20_00096670 | FBgn0026390 | DBIA002468 | augustus-dsubp_Seq360-processed-gene-13.56 |
| **Or35a** | DS20_00085320 | FBgn0028946 | DBIA006736 | maker-dsubp_Seq360-exonerate_est2genome-gene-52.17 |
| **Or42b** | DS20_00123340 | FBgn0033043 | DBIA015520 | maker-dsubp_Seq69-exonerate_protein2genome-gene-1.0 |
| **Or45a** | DS20_00129850 | FBgn0033404 | DBIA009648 | maker-dsubp_Seq361-exonerate_protein2genome-gene-64.61 |
| **Or45b** | DS20_00129670 | FBgn0033422 | DBIA009630 | maker-dsubp_Seq361-exonerate_protein2genome-gene-65.42 |
| **Or46a** | DS20_00135780 | FBgn0026388 | DBIA004133 | maker-dsubp_Seq361-exonerate_est2genome-gene-34.8 |
| **Or47a** | DS20_00137170 | FBgn0026386 | DBIA004259 | maker-dsubp_Seq361-exonerate_est2genome-gene-37.15 |
| **Or47b** | DS20_00136900 | FBgn0026385 | DBIA004235 | maker-dsubp_Seq361-augustus-gene-37.210 |
| **Or59a** | DS20_00133810 | FBgn0026384 | DBIA004638 | augustus-dsubp_Seq361-processed-gene-54.7 |
| **Or59c** | DS20_00133840 | FBgn0034866 | DBIA004642 | maker-dsubp_Seq12-exonerate_est2genome-gene-0.2 |
| **Or63a** | DS20_00003490 | FBgn0035382 | DBIA001054 | augustus-dsubp_Seq405-processed-gene-70.12 |
| **Or65c** | DS20_00005260 | FBgn0041623 | DBIA003777 | maker-dsubp_Seq405-snap-gene-77.414 |
| **Or67a** | DS20_00123090 | FBgn0036009 | DBIA011234 | augustus-dsubp_Seq355-processed-gene-14.24 |
| **Or67b** | DS20_00008700 | FBgn0036019 | DBIA000874 | maker-dsubp_Seq405-augustus-gene-33.226 |
| **Or67c** | DS20_00008100 | FBgn0036078 | DBIA000810 | maker-dsubp_Seq405-exonerate_est2genome-gene-89.3 |
| **Or67d** | DS20_00008080 | FBgn0036080 | DBIA000807 | maker-dsubp_Seq405-exonerate_est2genome-gene-89.12 |
| **Or69a** | DS20_00055820 | FBgn0041622 | DBIA003587 | maker-dsubp_Seq405-augustus-gene-61.185 |
| **Or7a** | DS20_00015570 | FBgn0030016 | DBIA010477 | maker-dsubp_Seq407-exonerate_est2genome-gene-35.1 |
| **Or85a** | DS20_00044650 | FBgn0037576 | DBIA001989 | maker-dsubp_Seq354-augustus-gene-37.324 |
| **Or85b** | DS20_00038870 | FBgn0037590 | DBIA002008 | maker-dsubp_Seq241-exonerate_est2genome-gene-0.3 |
| **Or85d** | DS20_00038820 | FBgn0037594 | DBIA002011 | maker-dsubp_Seq354-exonerate_protein2genome-gene-37.63 |
| **Or85e** | DS20_00038780 | FBgn0026399 | DBIA002017 | maker-dsubp_Seq354-exonerate_protein2genome-gene-37.117 |
| **Or85f** | DS20_00062320 | FBgn0037685 | DBIA002118 | maker-dsubp_Seq355-exonerate_protein2genome-gene-2.25 |
| **Or88a** | DS20_00053320 | FBgn0038203 | DBIA001455 | maker-dsubp_Seq354-exonerate_protein2genome-gene-22.15 |
| **Or94a** | DS20_00146650 | FBgn0039033 | DBIA001732 | augustus-dsubp_Seq355-processed-gene-35.12 |
| **Or98a** | DS20_00097740 | FBgn0039551 | DBIA009044 | maker-dsubp_Seq355-exonerate_est2genome-gene-24.7 |
| **Or98b** | DS20_00120440 | FBgn0039582 | DBIA005141 | augustus-dsubp_Seq355-processed-gene-20.1 |
| **Or9a** | DS20_00016110 | FBgn0030204 | DBIA007312 | snap-dsubp_Seq407-processed-gene-45.136 |
| **Gr10a** | DS20_00048010 | FBgn0045502 | DBIA005519 | maker-dsubp_Seq407-exonerate_protein2genome-gene-75.11 |
| **Gr10b** | DS20_00048000 | FBgn0030297 | DBIA005518 | maker-dsubp_Seq407-exonerate_protein2genome-gene-75.10 |
| **Gr21a** | DS20_00076230 | FBgn0041250 | DBIA008570 | augustus-dsubp_Seq14-processed-gene-1.49 |
| **Gr22a** | DS20_00083860 | FBgn0045501 | DBIA010841 | augustus-dsubp_Seq360-processed-gene-47.7 |
| **Gr22e** | DS20_00083850 | FBgn0045497 | DBIA010840 | maker-dsubp_Seq360-exonerate_protein2genome-gene-46.11 |
| **Gr28b** | DS20_00091740 | FBgn0045495 | DBIA003334 | augustus-dsubp_Seq360-processed-gene-29.49 |
| **Gr2a** | DS20_00050700 | FBgn0265139 | DBIA005880 | maker-dsubp_Seq407-augustus-gene-65.148 |
| **Gr33a** | DS20_00097080 | FBgn0032416 | DBIA002425 | maker-dsubp_Seq360-exonerate_protein2genome-gene-11.52 |
| **Gr36a** | DS20_00087360 | FBgn0045487 | DBIA003002 | maker-dsubp_Seq360-exonerate_protein2genome-gene-59.7 |
| **Gr39a** | DS20_00089860 | FBgn0264556 | DBIA002732 | augustus-dsubp_Seq360-processed-gene-36.9 |
| **Gr39b** | DS20_00089920 | FBgn0041245 | DBIA002724 | snap-dsubp_Seq360-processed-gene-35.21 |
| **Gr47b** | DS20_00136850 | FBgn0041241 | DBIA004230 | maker-dsubp_Seq361-exonerate_protein2genome-gene-36.86 |
| **Gr57a** | DS20_00136720 | FBgn0041240 | DBIA004215 | maker-dsubp_Seq361-exonerate_protein2genome-gene-36.33 |
| **Gr58a** | DS20_00128240 | FBgn0041239 | DBIA004802 | maker-dsubp_Seq361-snap-gene-69.168 |
| **Gr59a** | DS20_00129160 | FBgn0045483 | DBIA004706 | maker-dsubp_Seq361-exonerate_protein2genome-gene-67.12 |
| **Gr59f** | DS20_00133950 | FBgn0041234 | DBIA004654 | maker-dsubp_Seq361-exonerate_protein2genome-gene-54.22 |
| **Gr5a** | DS20_00102140 | FBgn0003747 | DBIA011874 | augustus-dsubp_Seq407-processed-gene-97.20 |
| **Gr61a** | DS20_00001190 | FBgn0035167 | DBIA010640 | maker-dsubp_Seq405-exonerate_est2genome-gene-51.9 |
| **Gr63a** | DS20_00074380 | FBgn0035468 | DBIA000047 | maker-dsubp_Seq405-augustus-gene-35.103 |
| **Gr64c** | DS20_00053760 | FBgn0045477 | DBIA000072 | maker-dsubp_Seq405-snap-gene-36.291 |
| **Gr66a** | DS20_00010480 | FBgn0035870 | DBIA001123 | maker-dsubp_Seq405-exonerate_protein2genome-gene-97.147 |
| **Gr68a** | DS20_00071190 | FBgn0041231 | DBIA012033 | maker-dsubp_Seq405-exonerate_protein2genome-gene-27.61 |
| **Gr77a** | DS20_00003900 | FBgn0045474 | DBIA003934 | maker-dsubp_Seq405-exonerate_protein2genome-gene-72.54 |
| **Gr85a** | DS20_00038310 | FBgn0045473 | DBIA001538 | maker-dsubp_Seq354-exonerate_protein2genome-gene-32.49 |
| **Gr89a** | DS20_00065090 | FBgn0038440 | DBIA011120 | maker-dsubp_Seq355-augustus-gene-11.147 |
| **Gr8a** | DS20_00105100 | FBgn0030108 | DBIA006013 | maker-dsubp_Seq407-snap-gene-84.248 |
| **Gr92a** | DS20_00019880 | FBgn0045471 | DBIA001362 | maker-dsubp_Seq354-exonerate_protein2genome-gene-18.12 |
| **Gr93c** | DS20_00000030 | FBgn0045469 | DBIA001929 | augustus-dsubp_Seq0-processed-gene-0.11 |
| **Gr93d** | DS20_00000040 | FBgn0045468 | DBIA001928 | maker-dsubp_Seq0-snap-gene-0.81 |
| **Gr94a** | DS20_00025130 | FBgn0041225 | DBIA001723 | maker-dsubp_Seq355-protein2genome-gene-35.105 |
| **Gr97a** | DS20_00041320 | FBgn0041224 | DBIA007734 | maker-dsubp_Seq355-exonerate_protein2genome-gene-55.32 |
| **Gr98a** | DS20_00097520 | FBgn0039520 | DBIA005065 | maker-dsubp_Seq355-exonerate_protein2genome-gene-23.13 |
| **Gr9a** | DS20_00104450 | FBgn0052693 | DBIA011665 | augustus-dsubp_Seq407-processed-gene-90.1 |
| **Ir100a** | DS20_00068470 | FBgn0039879 | DBIA014431 | maker-dsubp_Seq355-protein2genome-gene-66.170 |
| **Ir11a** | DS20_00111940 | FBgn0030385 | DBIA005011 | maker-dsubp_Seq407-exonerate_protein2genome-gene-24.22 |
| **Ir20a** | DS20_00120950 | FBgn0031181 | DBIA012954 | augustus-dsubp_Seq407-processed-gene-13.282 |
| **Ir48b** | DS20_00139560 | FBgn0033648 | DBIA006097 | maker-dsubp_Seq361-exonerate_protein2genome-gene-44.33 |
| **Ir48c** | DS20_00139610 | FBgn0033651 | DBIA006102 | maker-dsubp_Seq361-exonerate_protein2genome-gene-44.34 |
| **Ir52a** | DS20_00034570 | FBgn0034023 | DBIA006570 | snap-dsubp_Seq361-processed-gene-20.51 |
| **Ir56c** | DS20_00143920 | FBgn0034457 | DBIA010251 | maker-dsubp_Seq361-exonerate_protein2genome-gene-51.56 |
| **Ir56d** | DS20_00143930 | FBgn0034458 | DBIA010253 | maker-dsubp_Seq361-exonerate_protein2genome-gene-51.32 |
| **Ir60e** | DS20_00133340 | FBgn0035019 | DBIA004598 | maker-dsubp_Seq361-exonerate_protein2genome-gene-55.71 |
| **Ir67a** | DS20_00008790 | FBgn0036010 | DBIA000885 | maker-dsubp_Seq405-exonerate_protein2genome-gene-32.130 |
| **Ir67b** | DS20_00008060 | FBgn0036083 | DBIA000802 | maker-dsubp_Seq405-exonerate_protein2genome-gene-88.26 |
| **Ir68b** | DS20_00005790 | FBgn0036250 | DBIA000546 | maker-dsubp_Seq405-exonerate_protein2genome-gene-78.39 |
| **Ir7a** | DS20_00029670 | FBgn0029961 | DBIA013727 | maker-dsubp_Seq407-exonerate_protein2genome-gene-37.13 |
| **lr7b** | DS20_00029730 | FBgn0029965 | DBIA015424 | maker-dsubp_Seq407-exonerate_protein2genome-gene-36.42 |
| **lr7c** | DS20_00029750 | FBgn0029966 | DBIA015422 | maker-dsubp_Seq407-exonerate_est2genome-gene-36.10 |
| **lr7e** | DS20_00029770 | FBgn0259189 | DBIA015421 | maker-dsubp_Seq407-exonerate_protein2genome-gene-36.43 |
| **lr7g** | DS20_00029780 | FBgn0029968 | DBIA015420 | maker-dsubp_Seq407-exonerate_protein2genome-gene-36.58 |
| **lr85a** | DS20_00039730 | FBgn0037630 | DBIA002046 | augustus-dsubp_Seq355-processed-gene-0.17 |
| **lr87a** | DS20_00150770 | FBgn0038153 | DBIA001378 | maker-dsubp_Seq354-exonerate_protein2genome-gene-18.11 |
| **lr94e** | DS20_00123140 | FBgn0259194 | DBIA005346 | augustus-dsubp_Seq355-processed-gene-47.16 |
| **lr94g** | DS20_00123170 | FBgn0039079 | DBIA005349 | augustus-dsubp_Seq355-processed-gene-47.17 |
| **iav** | DS20_00049250 | FBgn0086693 | DBIA005662 | maker-dsubp_Seq407-exonerate_est2genome-gene-70.0 |
| **Mid1** | DS20_00133010 | FBgn0053988 | DBIA004559 | maker-dsubp_Seq361-exonerate_protein2genome-gene-56.37 |
| **Nach** | DS20_00130410 | FBgn0024319 | DBIA009703 | maker-dsubp_Seq361-snap-gene-63.184 |
| **nan** | DS20_00006980 | FBgn0036414 | DBIA000672 | maker-dsubp_Seq405-exonerate_est2genome-gene-83.0 |
| **nompC** | DS20_00077950 | FBgn0016920 | DBIA008361 | maker-dsubp_Seq360-exonerate_est2genome-gene-76.2 |
| **pain** | DS20_00127130 | FBgn0060296 | DBIA004909 | augustus-dsubp_Seq361-processed-gene-72.64 |
| **Piezo** | DS20_00093040 | FBgn0264953 | DBIA006910 | augustus-dsubp_Seq360-processed-gene-22.7 |
| **Pkd2** | DS20_00097250 | FBgn0041195 | DBIA002408 | augustus-dsubp_Seq360-processed-gene-13.18 |
| **ppk** | DS20_00084990 | FBgn0020258 | DBIA006781 | maker-dsubp_Seq360-exonerate_est2genome-gene-50.9 |
| **ppk11** | DS20_00092900 | FBgn0065109 | DBIA006927 | maker-dsubp_Seq360-exonerate_protein2genome-gene-22.40 |
| **ppk12** | DS20_00143270 | FBgn0034730 | DBIA010186 | maker-dsubp_Seq361-exonerate_est2genome-gene-29.6 |
| **ppk13** | DS20_00086820 | FBgn0053508 | DBIA003055 | maker-dsubp_Seq360-exonerate_protein2genome-gene-58.87 |
| **ppk15** | DS20_00101720 | FBgn0039424 | DBIA007678 | maker-dsubp_Seq355-exonerate_protein2genome-gene-57.9 |
| **ppk17** | DS20_00080850 | FBgn0032602 | DBIA012741 | maker-dsubp_Seq360-exonerate_protein2genome-gene-36.85 |
| **ppk18** | DS20_00092860 | FBgn0265001 | DBIA006928 | maker-dsubp_Seq360-snap-gene-22.88 |
| **ppk22** | DS20_00012680 | FBgn0051105 | DBIA001671 | maker-dsubp_Seq355-snap-gene-64.219 |
| **ppk23** | DS20_00051210 | FBgn0030844 | DBIA005824 | maker-dsubp_Seq407-exonerate_est2genome-gene-63.11 |
| **ppk25** | DS20_00033500 | FBgn0053349 | DBIA006456 | maker-dsubp_Seq361-snap-gene-17.700 |
| **ppk26** | DS20_00109310 | FBgn0035785 | DBIA004070 | maker-dsubp_Seq405-exonerate_protein2genome-gene-41.29 |
| **ppk28** | DS20_00113000 | FBgn0030795 | DBIA010422 | maker-dsubp_Seq407-exonerate_protein2genome-gene-53.9 |
| **ppk29** | DS20_00141090 | FBgn0034965 | DBIA009945 | maker-dsubp_Seq361-snap-gene-22.145 |
| **ppk31** | DS20_00074820 | FBgn0051065 | DBIA007762 | augustus-dsubp_Seq355-processed-gene-54.31 |
| **ppk5** | DS20_00149700 | FBgn0053289 | DBIA011492 | maker-dsubp_Seq405-exonerate_protein2genome-gene-17.40 |
| **ppk6** | DS20_00035060 | FBgn0034489 | DBIA008216 | augustus-dsubp_Seq361-processed-gene-48.26 |
| **ppk8** | DS20_00017250 | FBgn0052792 | DBIA007172 | maker-dsubp_Seq407-exonerate_protein2genome-gene-50.14 |
| **ppk9** | DS20_00128290 | FBgn0085398 | DBIA004798 | maker-dsubp_Seq361-augustus-gene-69.140 |
| **pyx** | DS20_00000480 | FBgn0035113 | DBIA010564 | maker-dsubp_Seq405-exonerate_protein2genome-gene-54.43 |
| **tmc** | DS20_00008720 | FBgn0267796 | DBIA000876 | maker-dsubp_Seq405-snap-gene-33.237 |
| **trp1** | DS20_00129450 | FBgn0005614 | DBIA009598 | maker-dsubp_Seq361-exonerate_est2genome-gene-66.2 |
| **Trpm** | DS20_00034790 | FBgn0265194 | DBIA009896 | maker-dsubp_Seq361-exonerate_est2genome-gene-21.0 |
| **Trpy** | DS20_00080660 | FBgn0032593 | DBIA012719 | maker-dsubp_Seq360-exonerate_est2genome-gene-36.7 |
| **wtrw** | DS20_00119270 | FBgn0260005 | DBIA008758 | augustus-dsubp_Seq354-processed-gene-6.71 |

**Table S7.** Sensory receptor expression patterns in abdominal tip. The expression (in Transcript Per Millions) of OR, GR, IR, and MR genes in the abdominal tip RNA-seq data for the genes that were under selection in the MK test in at least one species. Expressed genes are listed first. The most highly expressed sensory genes in the abdominal tip tissue are mechanosensory receptors. Most of OR, Gr and IR genes show little or no expression in the abdominal tip. TPMs are raw values for each species and are not normalized for samples between species. Expressed denotes genes showing TPM larger than 1 in at least one of the four species.

| Gene Name | *D. suzukii* | *D. subpulchrella* | *D. biarmipes* | *D. melanogaster* | Expressed |
| --- | --- | --- | --- | --- | --- |
| Or47a | 0.081 | 0.434 | 1.336 | 0.515 | Yes |
| Or85f | 0.062 | 23.286 | 3.624 | 0.599 | Yes |
| Gr39a | 0.196 | 0 | 1.152 | 1.012 | Yes |
| Gr63a | 1.978 | 7.836 | 0.429 | 0.112 | Yes |
| Gr64c | 0 | 0 | 5.273 | 0.057 | Yes |
| Ir87a | 1.086 | 0.151 | 0.058 | 0.155 | Yes |
| iav | 1.079 | 1.573 | 13.307 | 0.572 | Yes |
| Ir87a | 1.086 | 0.151 | 0.058 | 0.155 | Yes |
| nan | 0.029 | 0 | 1.457 | 0.252 | Yes |
| nompC | 0.705 | 0.569 | 0.678 | 1.3 | Yes |
| Piezo | 25.339 | 20.195 | 8.238 | 15.818 | Yes |
| Pkd2 | 0.152 | 0.476 | 16.248 | 0.042 | Yes |
| ppk | 1.976 | 2.278 | 4.353 | 2.365 | Yes |
| ppk13 | 0.052 | 0 | 6.399 | 0.151 | Yes |
| ppk17 | 13.025 | 21.831 | 12.007 | 0.957 | Yes |
| ppk22 | 0.152 | 0 | 19.193 | 0.082 | Yes |
| ppk26 | 2.987 | 4.15 | 4.452 | 2.313 | Yes |
| tmc | 0.158 | 0.114 | 1.287 | 1.108 | Yes |
| trp1 | 0.535 | 1.069 | 0.234 | 0.699 | Yes |
| trpm | 5.756 | 5.836 | 11.108 | 8.171 | Yes |
| wtrw | 19.143 | 21.61 | 26.798 | 9.313 | Yes |
| Or10a | 0.082 | 0.091 | 0.122 | 0 | No |
| Or13a | 0.107 | 0 | 0.556 | 0 | No |
| Or19b | 0.024 | 0.328 | 0.037 | 0 | No |
| Or1a | 0 | 0 | 0 | 0.485 | No |
| Or22a | 0.027 | 0 | 0.145 | 0.259 | No |
| Or2a | 0.526 | 0.57 | 0.31 | 0 | No |
| Or30a | 0 | 0 | 0 | 0 | No |
| Or33c | 0.097 | 0 | 0 | 0 | No |
| Or35a | 0 | 0.07 | 0 | 0 | No |
| Or45a | 0 | 0 | 0.155 | 0.013 | No |
| Or45b | 0 | 0 | 0.101 | 0 | No |
| Or46a | 0 | 0.163 | 0.11 | 0 | No |
| Or47b | 0 | 0 | 0 | 0.04 | No |
| Or63a | 0 | 0 | 0 | 0.027 | No |
| Or65c | 0 | 0 | 0 | 0.413 | No |
| Or67a | 0 | 0 | 0 | 0.033 | No |
| Or67b | 0.149 | 0 | 0.4 | 0 | No |
| Or67c | 0 | 0 | 0 | 0.04 | No |
| Or67d | 0 | 0.158 | 0 | 0 | No |
| Or69a | 0 | 0 | 0 | 0 | No |
| Or7a | 0 | 0.224 | 0.129 | 0.29 | No |
| Or85a | 0 | 0.525 | 0.385 | 0 | No |
| Or85b | 0 | 0 | 0.039 | 0 | No |
| Or85d | 0 | 0 | 0.097 | 0 | No |
| Or85e | 0 | 0 | 0 | 0.042 | No |
| Or88a | 0 | 0 | 0 | 0 | No |
| Or94a | 0 | 0.119 | 0.172 | 0.15 | No |
| Or98b | 0 | 0 | 0 | 0 | No |
| Or9a | 0 | 0.255 | 0 | 0.076 | No |
| Gr10a | 0 | 0 | 0 | 0.179 | No |
| Gr10b | 0 | 0 | 0.313 | 0.129 | No |
| Gr21a | 0.045 | 0 | 0.045 | 0.2 | No |
| Gr28b | 0.365 | 0.329 | 0.16 | 0.546 | No |
| Gr2a | 0.275 | 0.279 | 0 | 0.079 | No |
| Gr33a | 0.132 | 0 | 0 | 0.18 | No |
| Gr36a | 0 | 0 | 0.036 | 0 | No |
| Gr39b | 0.359 | 0.715 | 0 | 0.03 | No |
| Gr57a | 0 | 0 | 0 | 0 | No |
| Gr5a | 0.067 | 0.135 | 0.246 | 0.089 | No |
| Gr61a | 0.065 | 0.1 | 0 | 0.131 | No |
| Gr68a | 0 | 0 | 0 | 0.113 | No |
| Gr77a | 0.135 | 0.627 | 0 | 0.404 | No |
| Gr89a | 0 | 0 | 0.08 | 0.137 | No |
| Gr93c | 0 | 0 | 0.188 | 0 | No |
| Gr93d | 0 | 0 | 0.354 | 0.051 | No |
| Ir11a | 0 | 0 | 0.032 | 0 | No |
| Ir48b | 0 | 0.046 | 0.032 | 0.064 | No |
| Ir56c | 0 | 0 | 0.53 | 0 | No |
| Ir56d | 0.052 | 0 | 0 | 0 | No |
| Ir67a | 0 | 0 | 0 | 0.057 | No |
| Ir67b | 0 | 0.08 | 0.018 | 0 | No |
| **lr68b** | 0.073 | 0.066 | 0.235 | 0.061 | No |
| **lr7a** | 0 | 0 | 0 | 0 | No |
| **lr7c** | 0.051 | 0.065 | 0.113 | 0.12 | No |
| **lr7e** | 0 | 0 | 0.022 | 0 | No |
| **lr94e** | 0 | 0 | 0.063 | 0.058 | No |
| **Nach** | 0.229 | 0 | 0.066 | 0.073 | No |
| **pain** | 0.082 | 0.029 | 0.584 | 0.454 | No |
| **ppk12** | 0 | 0.122 | 0.106 | 0.147 | No |
| **ppk15** | 0 | 0 | 0.057 | 0.023 | No |
| **ppk23** | 0 | 0.141 | 0.275 | 0.324 | No |
| **ppk25** | 0.022 | 0 | 0.048 | 0.067 | No |
| **ppk29** | 0.188 | 0.078 | 0.128 | 0.333 | No |
| **ppk6** | 0 | 0 | 0.23 | 0.201 | No |
| **ppk8** | 0.043 | 0 | 0.633 | 0.032 | No |
| **ppk9** | 0.027 | 0 | 0 | 0.042 | No |
| **Trpy** | 0.082 | 0.029 | 0.584 | 0.454 | No |

